# Ancient and nonuniform loss of olfactory receptor expression renders the shark nose a *de facto* vomeronasal organ

**DOI:** 10.1101/2022.11.27.518102

**Authors:** Adnan S. Syed, Kanika Sharma, Maxime Policarpo, Sara Ferrando, Didier Casane, Sigrun I. Korsching

**Affiliations:** Institute of Genetics, Mathematical-Natural Sciences Faculty, University of Cologne, Germany; Institute of Virology, Faculty of Medicine and University Hospital of Cologne, University of Cologne, Germany; Université Paris-Saclay, CNRS, IRD, UMR Évolution, Génomes, Comportement et Écologie, 91190, Gif-sur-Yvette, France; Zoological Institute, Department of Environmental Sciences, University of Basel, Basel, Switzerland; Department of Earth, Environmental, and Life Sciences (DISTAV), University of Genoa, Italy; NBFC, National Biodiversity Future Center, Palermo 90133, Italy; Université Paris Cité, UFR Sciences du Vivant, F-75013 Paris, France

**Author notes:** participated equally in this work (Adnan S. Syed, Kanika Sharma, Maxime Policarpo, Sara Ferrando).

**Keywords:** gene family dynamics, gene expression, Chondrichthyes, odorant receptors, trace amine-associated receptors, vomeronasal receptors

## Abstract

Cartilaginous fishes are renowned for a keen sense of smell, a reputation based on behavioral observations and supported by the presence of large and morphologically complex olfactory organs. At the molecular level, genes belonging to the four families coding for most olfactory receptors in other vertebrates have been identified in a chimera and a shark, but it was unknown whether they actually code for olfactory receptors in these species. Here we describe the evolutionary dynamics of these gene families in cartilaginous fishes using genomes of a chimera, a skate, a sawfish and eight sharks. The number of putative OR, TAAR and V1R/ORA receptors is very low and stable whereas the number of putative V2R/OlfC receptors is higher and much more dynamic. In the catshark *Scyliorhinus canicula*, we show that many V2R/OlfC receptors are expressed in the olfactory epithelium in the sparsely distributed pattern characteristic for olfactory receptors. In contrast, the other three vertebrate olfactory receptor families are either not expressed (OR) or only represented with a single receptor (V1R/ORA and TAAR). The complete overlap of markers of microvillous olfactory sensory neurons with panneuronal marker HuC in the olfactory organ suggests the same cell type specificity of V2R/OlfC expression as for bony fishes, i.e. in microvillous neurons. The relatively low number of olfactory receptors in cartilaginous fishes compared to bony fishes could be the result of an ancient and constant selection in favor of a high olfactory sensitivity at the expense of a high discrimination capability.

## Introduction

The sense of smell is involved in many essential tasks of vertebrates, including Chondrichthyes (cartilaginous fishes), from food and prey location over reproductive functions and social interactions to danger avoidance (DeMaria et al. 2013; Gardiner et al. 2014; Gardiner et al. 2015). The study of Osteichthyes (bony fishes, such as mouse and zebrafish), but also lampreys (jawless fishes), has shown that four large olfactory receptor families (OR, TAAR, V1R/ORA and V2R/OlfC) are expressed in olfactory sensory neurons (OSNs) and constitute the molecular basis of odor detection (Mombaerts 2004). These families of olfactory receptors were first identified in mammals (Buck and Axel 1991; Dulac and Axel 1995; Matsunami and Buck 1997; Liberles and Buck 2006), but subsequent studies have shown their presence in other tetrapods and Actinopterygii (ray-finned fishes) (Niimura 2009), in cartilaginous fishes (Grus and Zhang 2009; Hussain et al. 2009; Niimura 2009; Sharma et al. 2019) and in jawless fishes (Grus and Zhang 2009; Libants et al. 2009; Dieris et al. 2021; Kowatschew and Korsching 2022), suggesting that they were present in the last common ancestor of all extant vertebrates.

We have recently described the olfactory repertoire of the small-spotted catshark *Scyliorhinus canicula* to be dominated by the VR2/OlfC family, whereas the VR1/ORA, OR and TAAR families are only represented by a handful of members each (Sharma et al. 2019). This repertoire is similar to that of four other sharks - the cloudy catshark *Scyliorhinus torazame*, the brownbanded bamboo shark *Chiloscyllium punctatum*, the whale shark *Rhincodon typus* and the white shark *Carcharodon carcharias* (Hara et al. 2018; Marra et al. 2019) - and a more distantly related species, the chimaera *Callorhinchus milii* (Grus and Zhang 2009; Hussain et al. 2009; Niimura 2009). These olfactory gene repertoires are distinctly different from those of both jawless and bony fishes, which are dominated by the OR family (Niimura 2009).

The expression of olfactory receptor genes has been overwhelmingly studied in mammals, in particular mouse and rat, and to some extent in teleosts, in particular zebrafish. The expression for all four families is very similar in mammals and teleosts: a monogenic expression pattern and a characteristically sparse expression of individual receptor genes are features common to both (Mombaerts 2004; Korsching 2020a). In both mammals and teleosts, olfactory receptor expression is segregated between two main types of OSNs, ciliated and microvillous neurons. ORs and TAARs are expressed in ciliated neurons, whereas V2R (in mammals also V1R) are expressed in microvillous neurons (Korsching 2020b). Much less is known about the expression of olfactory receptors in jawless fishes (Berghard and Dryer 1998; Freitag et al. 1999; Libants et al. 2009; Kowatschew and Korsching 2022). To the best of our knowledge, no *in situ* expression studies have been performed for any olfactory receptor of any cartilaginous fish (sharks, rays and chimaeras).

The olfactory organ of cartilaginous fishes looks similar to that of many ray-finned fishes: a rosette with bilaterally symmetric rows of lamellae (fig. 1). However, cartilaginous fishes always exhibit so-called secondary lamellae, emanating from the primary lamella, which also contain olfactory epithelium (fig. 1). Furthermore, and in contrast to tetrapods and ray-finned fishes, cartilaginous fishes seem not to possess ciliated OSNs (Holl 1973; Theisen et al. 1986; Takami et al. 1994) and their OSN repertoire is dominated by microvillous neurons, with rare crypt neurons (Holl 1973; Theisen et al. 1986; Takami et al. 1994; Ferrando et al. 2006). Thus, it is an open question, if and how ORs and TAARs - which are expressed in ciliated OSNs in both ray-finned fishes and tetrapods - are expressed in cartilaginous fish OSNs.

**Fig. 1.**
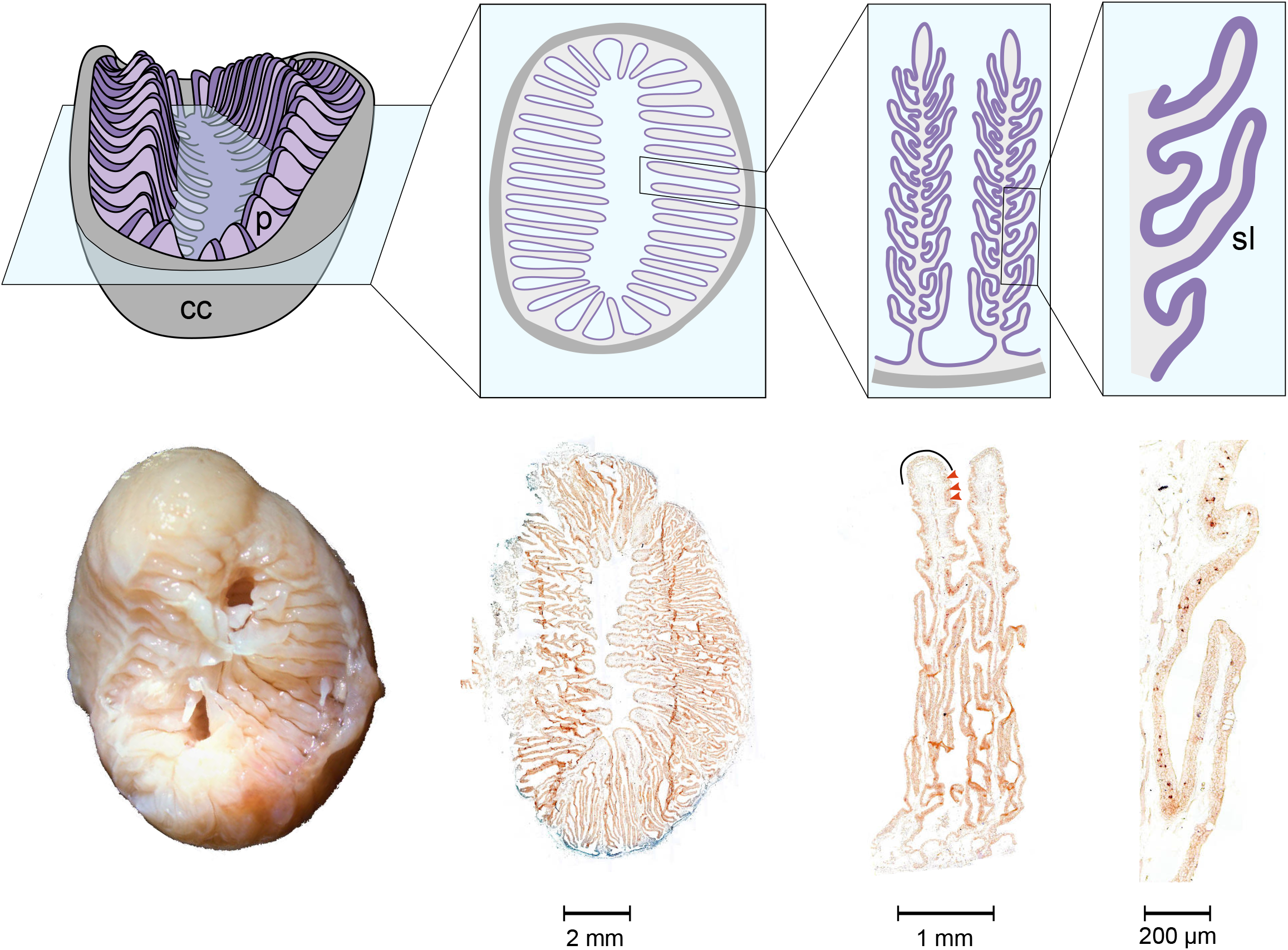
Morphology of the catshark olfactory organ. The catshark olfactory organ is shown at increasing resolution from left to right, starting with the whole organ enclosed by a connective capsule (cc), down to a segment of a single lamella. Top row, schematic representation; bottom row, micrographs of the same features, scale bars as indicated. Micrograph sections are from an *in situ* hybridisation experiment. Note the presence of primary (p) and secondary (sl) lamellae. The sensory surface covers both primary and secondary lamellae, excluding lamellar tips (solid black line), where mucous cells (large ovals) are enriched. Some labelled OSNs are pointed out by arrowheads (brown).

Here we made use of the recent availability of genomes for species from the three main groups of cartilaginous fishes - chimaeras, rays/skates/sawfishes and sharks, in total eleven species - to obtain a comprehensive picture of the evolutionary dynamics of the four olfactory gene families in chondrichthyans. We report that consistently OR, TAAR and V1R/ORA repertoires are very small and stable, whereas the V2R/OlfC repertoire is larger and more dynamic.

Furthermore, we examined the expression of olfactory genes in the catshark *S. canicula*. None of the few *or* genes present in its genome are expressed in the olfactory organ. For TAAR and V1R/ORA families, we observed expression in the olfactory organ for a single gene each. In contrast, VR2/OlfCs showed robust expression with several different probes. Individual olfactory receptor genes are expressed in sparsely distributed cells, and their spatial expression patterns are characteristically different between different receptors, both features as observed in other vertebrates. Globally, these results suggest that olfaction in cartilaginous fishes essentially relies on a relatively small set of *v2r/olfC* genes and that in several aspects the olfactory system of this vertebrate class could be considered a vomeronasal system (*cf*. Ferrando and Gallus, 2013).

## Results

### Diversity and evolutionary dynamics of the olfactory gene repertoire in cartilaginous fishes

We counted the number of complete coding sequences, pseudogenes, truncated and edge belonging to the OR, TAAR, V1R/ORA and V2R/OlfC families in eleven genomes of cartilaginous fishes. Five genomes have been previously examined but often only the number of complete genes has been reported and not always for all gene families. For *S. canicula* we refined the identification of complete genes in each family (*cf*. Sharma et al., 2019). For *C. milii*, the number of complete *or* and *v2r/olfC* genes were underestimated. For *S. torazame, C. punctatum* and *R. typus*, the number of complete genes were much underestimated or not examined at all. Once corrected, the numbers of genes in these species were similar and similar to those found in the six species for which the olfactory gene repertoire has never been studied before (fig. 2, supplementary fig. S1, supplementary table S1, Supplementary Material online).

**Fig. 2.**
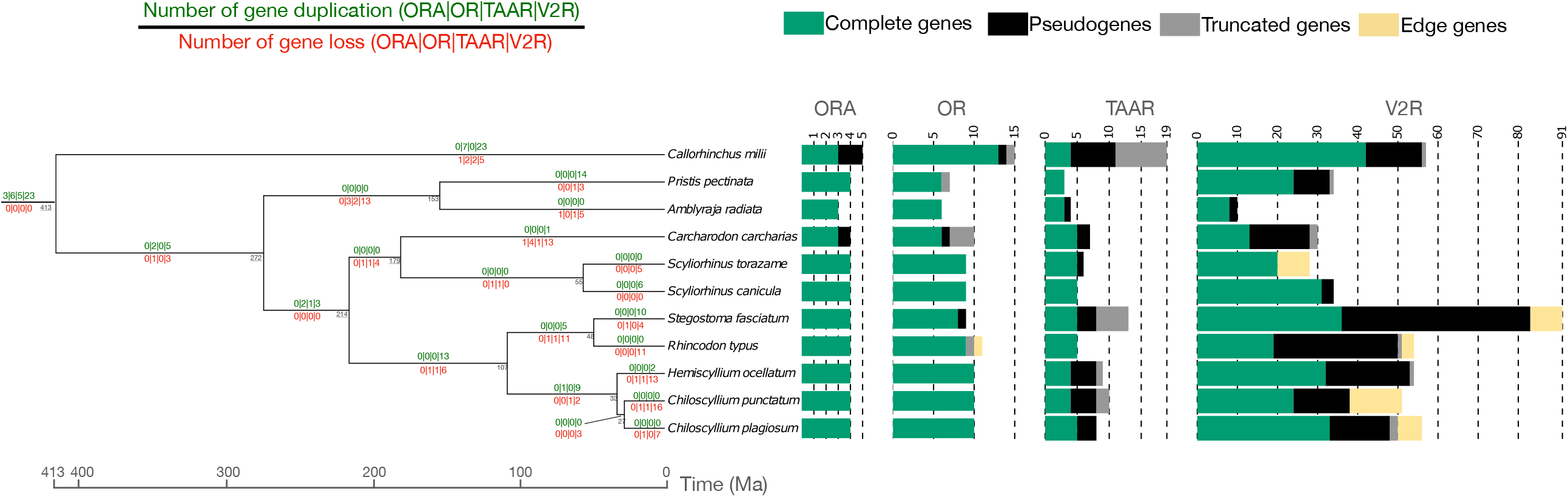
Time-calibrated chondrichthyan tree. The species tree topology was inferred by maximum likelihood using 1068 BUSCO genes and node ages were inferred using the least square dating method. The number of genes in the four olfactory receptor families are represented by multiple values barplots. Number of gene losses and gene gains in each branch of the tree and for the four gene families were inferred using the gene tree – species tree reconciliation method. The complete species tree with confidence intervals of node dates and with *Hydrolagus affinis* is available in supplementary fig. S5, Supplementary Material online.

In the cartilaginous fishes examined, the number of *or* genes varies between 6 and 13, and only 2 pseudogenes, 7 truncated and 1 edge sequences were found. This pattern implies 12 gene duplications and 18 gene losses, and the presence of seven *or* genes in the last common ancestor. The number of *taar* genes varies between 3 and 5, and in total 25 pseudogenes, and 15 truncated sequences were found. This pattern implies 1 gene duplication and 14 gene losses, and the presence of six *taar* genes in the last common ancestor. The number of *v1r/ora* genes varies between 2 and 4, and only 3 pseudogenes were found. This pattern implies no gene duplication and 6 gene losses and the presence of four *ora* genes in the last common ancestor. The number of *v2r/olfC* genes varies between 8 and 43, and 189 pseudogenes, 16 truncated and 15 edge sequences were found. This pattern implies 94 gene duplications and 121 gene losses, and the presence of twenty-one *v2r/olfC* genes in the last common ancestor (fig. 2).

Overall, it appears that the number of *or, taar* and *v1r/ora* genes is low and stable and very few pseudogenes and truncated genes are present in the genome of cartilaginous fishes, in deep contrast with large and highly dynamic numbers of genes belonging to these gene families in bony fishes, often associated with the presence of many pseudogenes and truncated genes. On the contrary, and as in bony fishes, the number of *v2r/olfC* is often high and much more variable. Several species-specific expansions exist (supplementary fig. S1, supplementary table S1, Supplementary Material online), and large proportion of pseudogenes are often present, a hallmark of a multigene family coding for olfactory receptors.

### Microvillous neuronal markers TRPC2 and Go label the entire neuronal population in the catshark olfactory epithelium

We employed the pan-neuronal marker HuC to visualize the entire neuronal population in the catshark olfactory epithelium (fig. 3). The lamellae of the olfactory organ are covered almost entirely by the sensory olfactory epithelium; only the tip region of the lamellae is covered by non-sensory epithelium (fig. 1). The HuC-immunoreactive neurons form an almost continuous irregular monolayer of pericarya (fig. 3b), which are situated in the middle layer of the epithelium, below the apical layer of supporting cells recognizable by their palisade-like arrangement. Proliferating cells form the basal layer of the olfactory epithelium and were visualized by proliferating cell nuclear antigen (PCNA) antibody (fig. 3). As expected, no overlap between HuC and PCNA immunoreactivity was observed (fig. 3a, d).

**Fig. 3.**
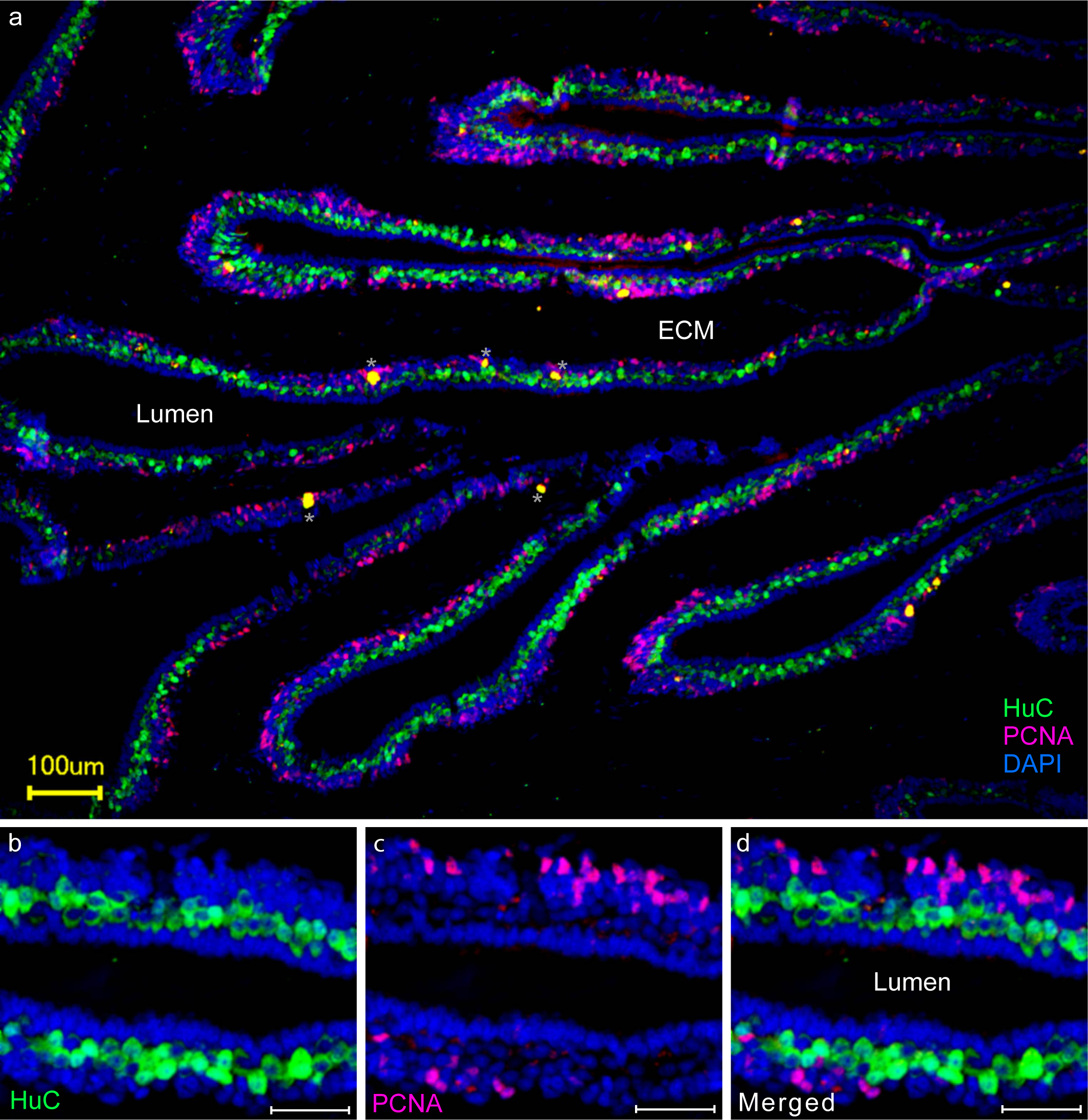
Catshark OSNs form a monolayer above the proliferative zone of the olfactory epithelium. Double immunofluorescence for a pan-neuronal marker (HuC, green) and a marker for mitotic cells (proliferating cell nuclear antigen, PCNA, purple) was performed on cryostat sections of the olfactory organ of the catshark. Nuclei are stained by DAPI (blue). Neurons (green) form a irregular monolayer below the supporting cells (dense palisade facing the lumen, blue) and above the basal layer (purple cells). a) overview, merged fluorescence, b-d) higher magnification, fluorescent label as indicated. Scale bars, 100 μm for panel a) and 40 μm for panels b-d). Asterisks, no nuclei are associated with these structures. ECM, extracellular matrix.

We then examined the expression of two established microvillous markers (transient receptor potential channel TRPC2 and G alpha protein Go (Hansen et al. 2004; Sato et al. 2005) within the entire neuronal population as defined by HuC immunoreactivity. Notwithstanding different subcellular compartments for HuC (perikarya) and Go (dendrites and axons), all HuC-positive cells appear to be Go-positive (fig. 4d, g). This was confirmed in double-labeling experiments using HuC antibody and Go *in situ* hybridisation (fig. 4h-j). Moreover, TRPC2, which labels all microvillous neurons in bony fishes (Sato et al. 2005; Omura and Mombaerts 2014), co-localizes completely with HuC immunoreactivity (fig. 4k-m). We did not detect any HuC-positive but TRPC2-negative or Go-negative cells. However, we cannot exclude the potential presence of a minor population of Go-negative ciliated neurons, on the scale of crypt neuron frequency (i.e., very minor), since this population is known to be negative for Go (Ferrando et al. 2009) and was not detected in our analysis. Nevertheless, the present data are consistent with the absence of ciliated OSNs in catshark olfactory epithelium.

**Fig. 4.**
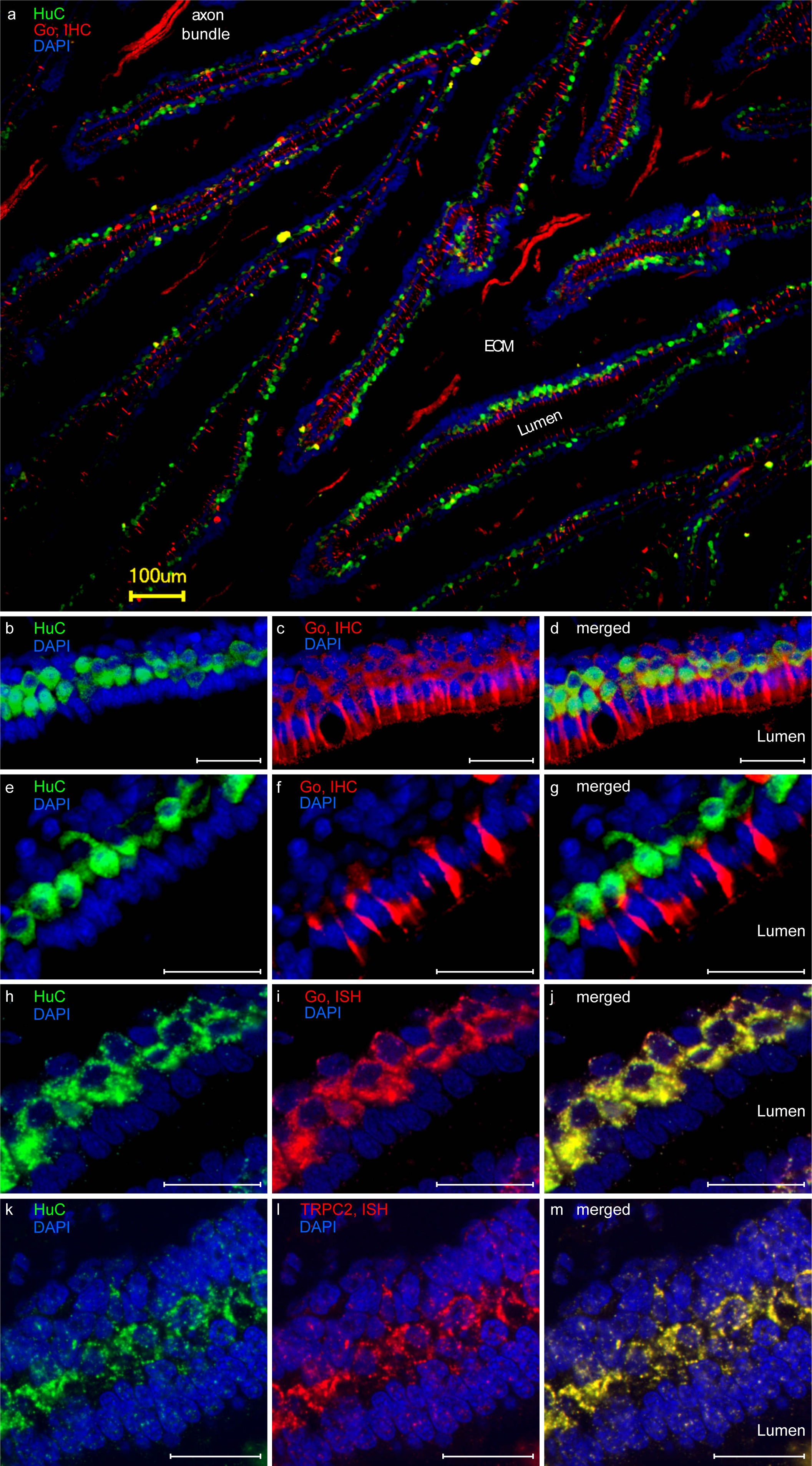
Go and TRPC2 label the entire OSN population. Cryostat sections of the olfactory organ of the catshark. ECM, extracellular matrix. a-g). Double immunofluorescence for Go (red) and HuC (green), nuclei are stained by DAPI (blue). a, c, f) Go immunoreactivity is seen in dendrites and axon bundles in the lamina propria of the lamellae, labelled here as ECM(extracellular matrix). Single axons are below threshold. d) Note that due to different subcellular localization the overlap between HuC and Go immunoreactivity, albeit clearly visible, is limited to the base of the dendrites. h-j) HuC immunofluorescence (green) and *in situ* hybridisation for Go (red). All HuC-immunoreactive cells express Go. k-m) HuC immunofluorescence (green) and *in situ* hybridisation for TRPC2 (red). All HuC-immunoreactive cells are labelled with the probe for TRPC2. Scale bars, 100 μm for panel a) and 40 μm for panels b-m).

Taken together, (nearly) all OSNs within the sensory surface of the catshark olfactory epithelium appear to express Go and TRPC2, suggesting that the entire OSN population of catshark consists of microvillous neurons. This fits well with the predominance of V2R/OlfC in the olfactory receptor repertoire of cartilaginous fishes, since in both tetrapods and teleosts V2R/OlfC receptors are characteristically expressed in microvillous neurons, and absent from ciliated neurons (Hansen et al. 2004; Mombaerts 2004; Syed et al. 2017).

### A comprehensive approach to study expression of the entire olfactory repertoire

The catshark olfactory receptor repertoire is dominated by V2Rs/OlfCs, with 34 *v2r/olfC* genes in contrast to 4-9 receptors for the other three families (OR, TAAR, V1R/ORA) (fig. 2, supplementary table 1, Supplementary Material online). Here we have examined the expression patterns for all four olfactory receptor families in catshark.

We performed RT-PCR for all *or, taar, taar-like* and *v1r/ora* genes identified in (Sharma et al., 2019) and for one *v2r/olfC-like* and five *v2r/olfC* genes (*v2rl4, v2r1, v2r6, v2r14, v2r19*, and *v2r29*). With exception of *v2rl4*, expression was observed for all genes examined (supplementary fig. S2, Supplementary Material online).

To examine expression at the cellular level, we performed *in situ* hybridisation with cRNA probes on horizontal cryostat sections from adult catshark olfactory epithelia (fig. 1). For the V2R/OlfC gene family, we used both specific and cross-reacting probes. The expression of *v2r1* and two *v2r*-like genes (*v2rl1, v2rl3*) was analysed with specific probes. In addition, we employed four probes from different *v2r* subclades which are expected to crossreact with several other genes (table 1), resulting in coverage of a considerable proportion of that family. For three families, ORs, TAARs, and V1R/ORAs, we examined the expression for each gene with a single, specific probe. In all cases where the characteristic pattern of sparsely distributed labelled cells was observed (supplementary fig. S3, Supplementary Material online), we quantified the expression frequency as well as the spatial distribution of receptor-expressing cells.

**Table 1.** Quantitative evaluation of expression for olfactory receptor genes from three different families.

| 1A |  |  |  |  |  |  |
| --- | --- | --- | --- | --- | --- | --- |
| Olfactory receptor |  | Density<br>(# OSN / mm lamellar<br>length) | % neurons in<br>primary/secondary<br>lamellae |  |  | # cross-reacting<br>genes |
| *ora2* |  | 0.45 +/- 0.05 | 47.3 / 52.7 |  |  | 0 |
| *taar1a* |  | 2.76 +/- 0.24 | 27.5 / 72.5 |  |  | 0 |
| *v2r1* |  | 5.94 +/- 0.81 | 81 / 19 |  |  | 0 |
| *v2r6* |  | 0.89 +/- 0.05 | 62.9 / 37.1 |  |  | 6 |
| *v2r14* |  | 1.075 +/- 0.06 | 47.3 / 52.7 |  |  | 1 |
| *v2r19* |  | 0.71 +/- 0.04 | 78.9 / 1.1 |  |  | 3 |
| *v2r29* |  | 0.78 +/- 0.07 | 85.8 / 14.2 |  |  | 1 |
| 1B |  |  |  |  |  |  |
| p values | *ora2* | *taar1a* | *v2r1* | *v2r6* | *v2r14* | *v2r19* |
| *ora2* |  | <0.01 | <0.001 | n.s | <0.001 | <0.001 |
| *taar1a* | <0.001 |  | <0.001 | <0.001 | <0.001 | <0.001 |
| *v2r1* | <0.001 | <0.001 |  | <0.05 | n.s | n.s |
| *v2r6* | <0.001 | <0.001 | <0.001 |  | <0.05 | <0.01 |
| *v2r14* | <0.001 | <0.001 | <0.001 | <0.001 |  | n.s |
| *v2r19* | <0.01 | <0.001 | <0.001 | <0.05 | <0.001 |  |
| *v2r29* | <0.001 | <0.001 | <0.001 | n.s | <0.01 | n.s |

**1A**, Density of OSN expressing a particular olfactory receptor gene is given as number of labelled OSNs per mm lamellar length. Distribution of labelled OSN between primary and secondary lamellae is given as percentage of total cells. Values are given as mean +/- SEM (110≤n<410). **1B**, p-values are estimated by t-test (two-sided, non-paired) and shown as matrix. Top triangle, p values for primary/secondary lamellae distribution; bottom triangle, p values for density of OSN comparison.

### The earliest-diverging V2R/OlfC gene exhibits the highest frequency of expression

A common feature of the tetrapod and teleost V2R/OlfC repertoires is the presence of genes coding for V2R/OlfC co-receptors and belonging to a monophyletic sister group of the main group of V2R/OlfC receptors. This receptor is a single gene in zebrafish, *olfCc1* (DeMaria et al. 2013), but has expanded to a small family in rodents, *vmn2r*, also known as *v2r2* (Martini et al. 2001). Because these genes serve as co-receptor for many individual *v2r/olfC* genes (Ishii and Mombaerts 2011; Akiyoshi et al. 2018), their expression frequency is characteristically high compared to the other *v2r/olfC* genes (DeMaria et al. 2013). The V2R1 receptor of catshark is the ortholog of OlfCc1 and Vmnr2r1-7. We were therefore interested in determining its expression frequency in relation to that of other *v2r/olfC* genes.

We performed *in situ* hybridisation for *v2r1*, two of the five *v2r/olfC-like* (*v2rl1,3*) and four *v2r* from the main clade *v2r6, v2r14, v2r19, v2r29*, which are expected to cross-react with one to six other v2r/olfC (see table 1 for details). No expression in the OE was seen for the *v2r/olfC-like* genes, but all other probes resulted in labelling of sparse cells within the OE as expected for olfactory receptor genes (figs. 5, 6). The gene *v2r1* was expressed in a considerable population of neurons (fig. 5 a-c), which appeared to be clearly larger than the populations labelled by each of the 4 cross-reacting probes (fig. 6, table 1). For a quantitative evaluation of position see below.

**Fig. 5.**
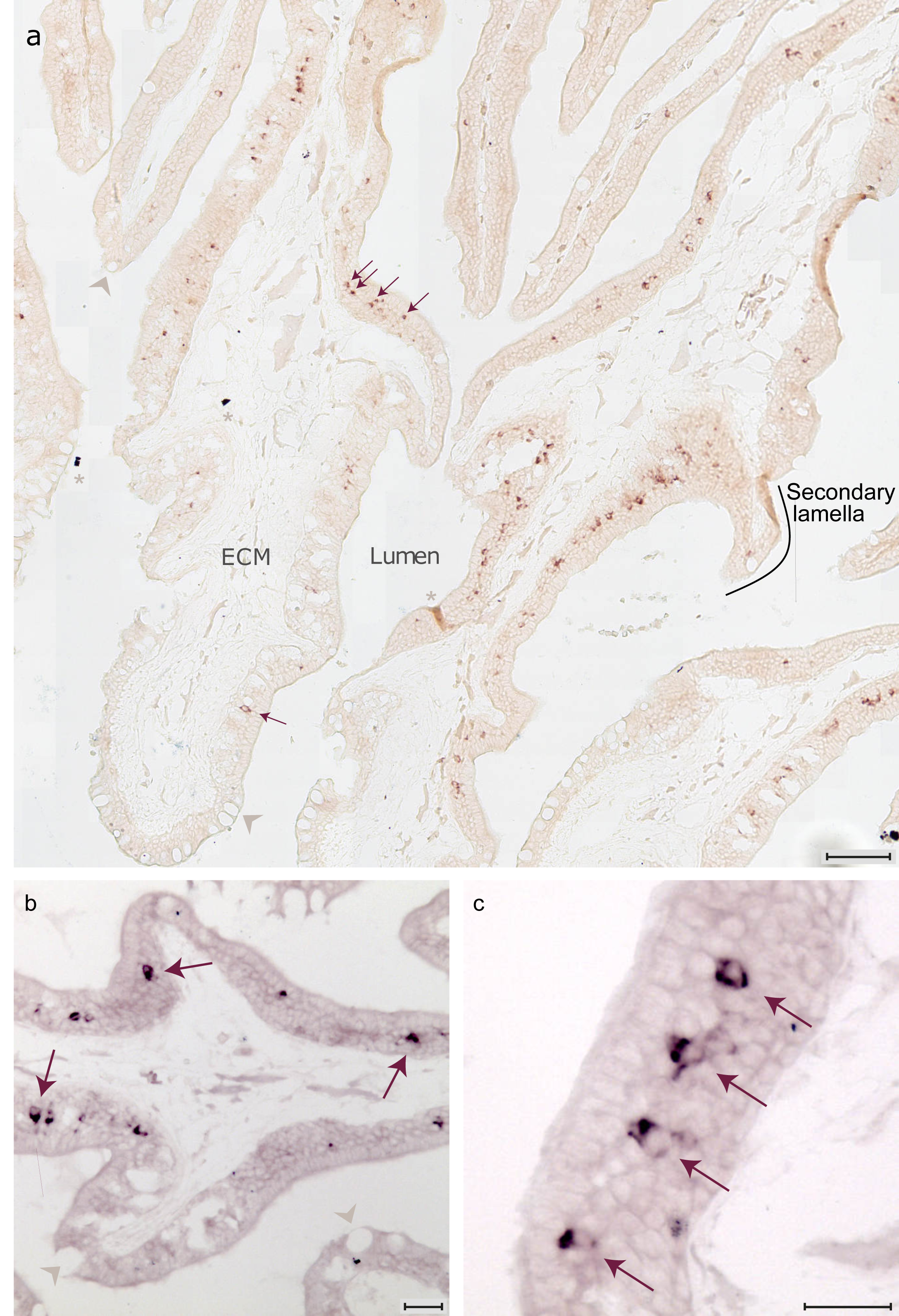
The basal gene of the main *v2r* clade, *v2r1*, is expressed at moderate frequency in the OE. Horizontal cryostat sections of the olfactory organ of the catshark were hybridised with a probe for *v2r1*. (a) *v2r1*-expressing OSNs are localized in the middle layer of the sensory epithelium, along both the primary and the secondary lamellae. A secondary lamella is indicated (black line). ECM, extracellular matrix; asterisks, artifacts; grey arrowheads, mucous cells. Scale bar, 100 μm. (b-c) Higher magnifications from different sections, some labeled neurons are pointed out by magenta arrows. Scale bar, 40 μm.

**Fig. 6.**
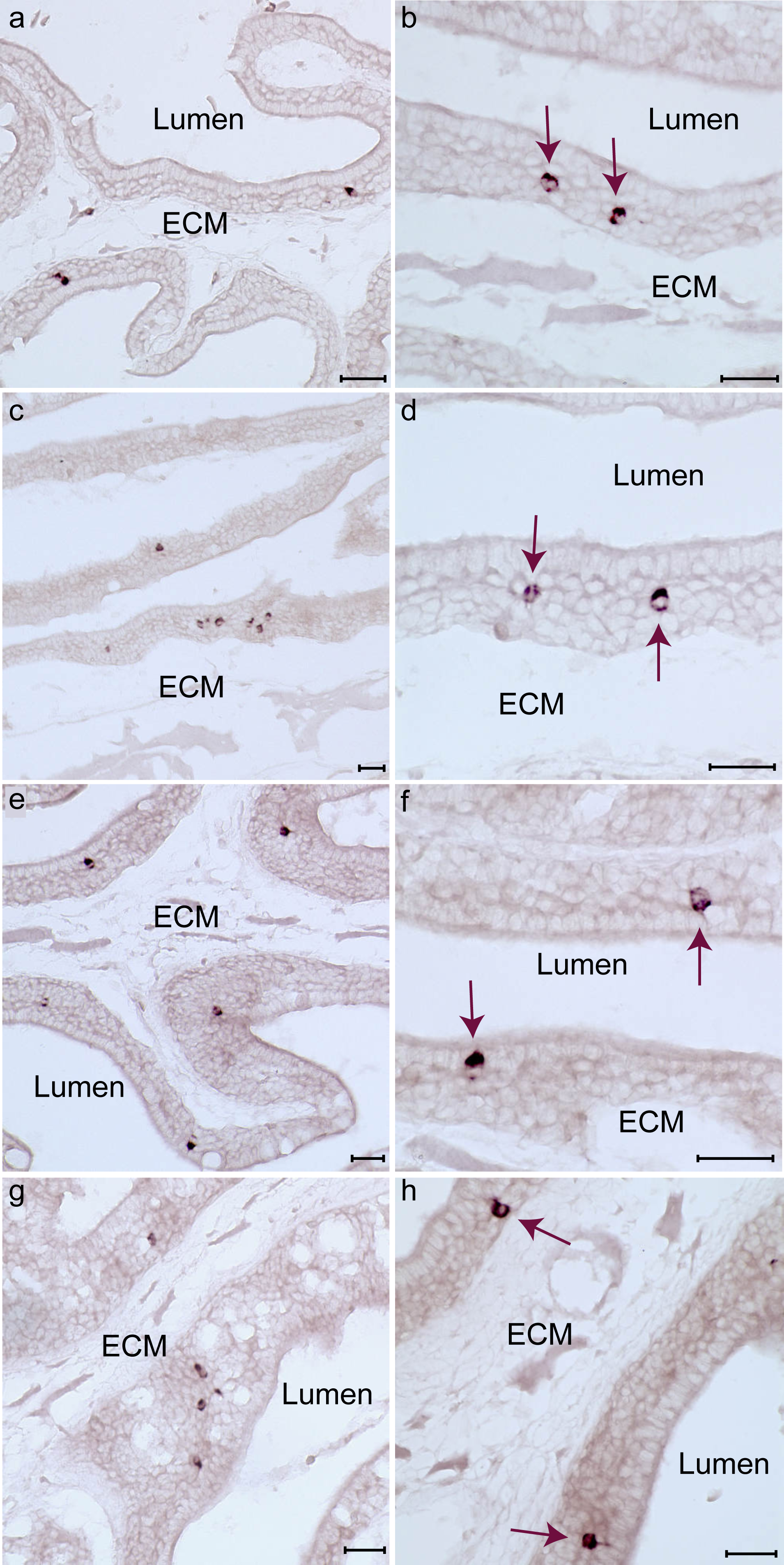
Sparse expression of *v2r* genes belonging to the main clade. Horizontal cryostat sections of catshark olfactory epithelium were hybridized with probes for *v2r29* (a,b), *v2r19* (c,d), *v2r6* (e,f), and *v2r14* (g,h). Right column, higher magnifications; all scale bars represent 40 μm. All probes show expression in small subsets of scattered OSNs, which are situated on primary and secondary lamellae. ECM, extracellular matrix. Some labelled neurons are indicated by magenta arrows.

### Two of the three minor olfactory receptor families are expressed in catshark OSNs

We analysed the expression of all *or, taar* and *v1r/ora* genes in the catshark olfactory receptor repertoire (Sharma et al. 2019) by *in situ* hybridisation. All probes were generated from olfactory organ RNA, allowing a first glimpse at expression. Indeed, we identified expression of all *or, taar* and *v1r/ora* genes in the RT-PCR of olfactory organ of catshark (supplementary fig. S2, Supplementary Material online).

In cartilaginous fishes, the OR family is very small and the OSN normally expressing ORs are absent (Holl 1973; Theisen et al. 1986; Takami et al. 1994; Hara et al. 2018; Marra et al. 2019; Sharma et al. 2019). Here we examined the expression of all catshark *ors* identified by (Sharma et al. 2019) with individually specific probes using *in situ* hybridisation. In no case expression could be seen, suggesting that the levels of mRNA for *ors* are sufficient for the more sensitive RT-PCR, but below the detection threshold of *in situ* hybridisation.

The V1R/ORA family of cartilaginous fish is small, similar to that of many teleosts (Saraiva and Korsching 2007; Zapilko and Korsching 2016), but see (Policarpo et al., 2022). All genes were examined for expression individually using *in situ* hybridisation. Expression was observed only for *ora2*. Very sparse cells situated in the neuronal layer (midlayer) of the olfactory lamellae are labeled (supplementary fig. S4a-d, Supplementary Material online, table 1).

The TAAR family of catshark consists of three *taar* genes proper and two *taar-like* (*tarl*) genes. *In situ* hybridisation with individually specific probes showed expression of taar1a in sparse neurons within the sensory surface of the olfactory organ (supplementary fig. S4e-h, Supplementary Material online), with clearly higher expression frequency than that observed for *ora2* (table 1). No expression was seen for the other two *taar* genes and the two *tarl* genes. The latter parallels the non-olfactory expression of *tarl* genes in bony fishes (Dieris et al. 2021).

### Distinctly different spatial distributions of neurons expressing different olfactory receptor genes

A characteristic feature of olfactory receptor expression in vertebrates is the restriction of expression of individual receptor genes to so-called expression zones or domains. Here we wished to investigate, whether similar patterns are present in a shark olfactory epithelium. Moreover, we examined whether there are differences between primary and secondary lamellae in terms of receptor expression.

We report that the ratio of expression (primary to secondary lamellae) is significantly different between receptor families (fig. 7b, table 1). *v2r1*-expressing cells are predominantly located on primary lamellae, whereas *ora2/v1r2*-expressing neurons have an equal probability to be present in primary and secondary lamellae. *taar1a*-expressing neurons show a third type of distribution and are strongly enriched on the secondary lamellae (fig. 7). Moreover, within the V2R/OlfC family, individual genes show different distributions, with *v2r6*-expressing cells showing the smallest preference for primary lamellae (fig. 7b, table 1).

**Fig. 7.**
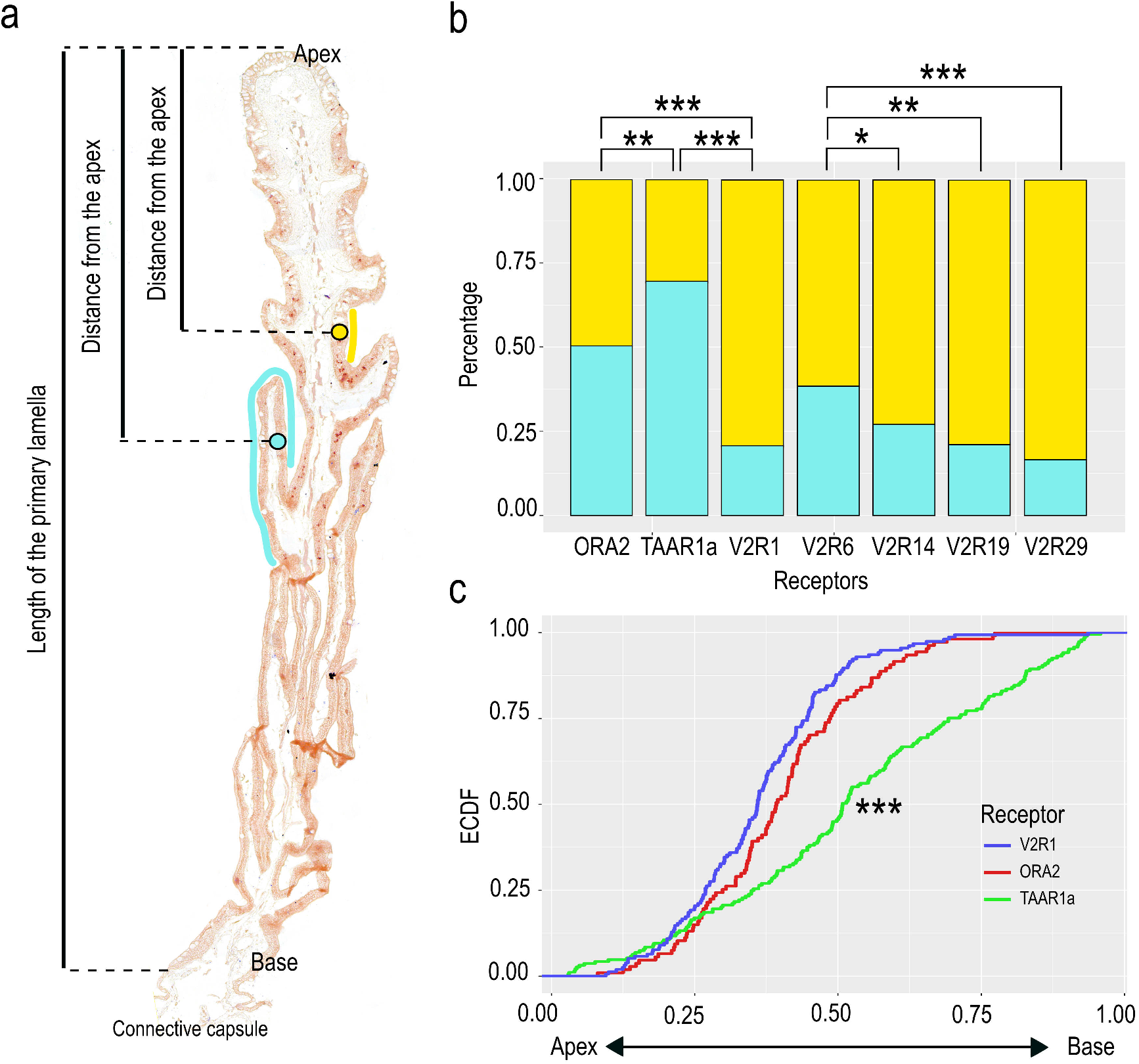
*Taar, ora*, and *v2r* genes show distinctly different, if overlapping spatial patterns of expression. a) Micrograph of a primary lamella with secondary lamellae from the olfactory organ of a catshark. To analyse the distribution of neurons expressing particular genes, the distance from the apex was measured and normalized to the total lamellar length from apex to base, both for neurons situated on primary and secondary lamellae (yellow and cyan circles, respectively). b) Percentage of neurons located on primary or secondary lamellae is shown as bar graph for seven genes from three different olfactory receptor families as indicated. Asterisks denote significance, see table 1 for numerical values. c) For three genes the position of labeled neurons along the lamellar length (cf. panel a) is shown as empirical cumulative distribution function (ECDF); blue, *v2r1*, red, *ora2*, green, *taar1a*. Significance of observed differences in distributions was estimated by Kolmogorov-Smirnov test (Marsaglia et al., 2003). ***, p<0.001 for *taar1a vs*. each of the other two genes. Distributions for *v2r1* and *ora2* are not significantly different from each other (see table 1 for exact values).

In bony vertebrates the average distance of OSNs from the center of the olfactory epithelium (radial distance) is characteristically different for different olfactory receptors (Weth et al. 1996; Mombaerts 2006). A recent study showed similar differences in radial distribution in a jawless fish (Kowatschew and Korsching 2022). Here, we have evaluated a similar parameter, the distance of labeled neurons from the base of the lamella, close to the connective capsule, which can also be considered a radial parameter. The distance was normalized to total length of the primary lamella (for details of the measurements, see Materials and Methods). We observed very similar distributions for *v2r1*- and *ora2*-expressing neurons, but a highly significant difference to the distribution of *taar1a*-expressing cells (fig. 7). Over three quarters of the *v2r1*- and *ora2*-expressing neurons are localized in the apical half of the lamellae, whereas the *taar1a*-expressing neurons are rather homogenously distributed along the baso-apical axis (fig. 7c).

Overall, considering the distribution between primary and secondary lamellae together with the radial distribution (along the lamellar axis) we found several distinctly and significantly different spatial patterns among the genes investigated. While the differences in basic morphology between cartilaginous, jawless and ray-finned fishes do not lend themselves to a direct comparison between spatial patterns, we wish to emphasize that the underlying principle of non-random spatial organization is shared among all three groups and thus may have emerged in the common ancestor of extant vertebrates.

## Discussion

### Evolutionary dynamics of the olfactory gene repertoire in cartilaginous fishes

We present a comprehensive analysis of the evolutionary dynamics of gene families coding for putative olfactory receptors in cartilaginous fishes, using eleven genomes of species belonging to the three main clades, *i*.*e*. chimaeras, rays/skates/sawfishes and sharks. The patterns of family size variation are strikingly different to those observed in bony fishes, the other clade of jawed vertebrates. First of all, the number of olfactory coding genes is on average much smaller and more stable in cartilaginous fishes. In most bony fishes, there are hundreds to thousands of olfactory receptor coding genes, but the genome of cartilaginous fishes codes only for approximatively 10 to 50 olfactory receptors. Such a low number of olfactory receptors is otherwise only known for species having a highly degenerated olfactory system, e.g. toothed whales (Kishida et al. 2015) and ocean sunfishes (Policarpo et al. 2021, 2022). However, in contrast to these species, cartilaginous fishes have well developed olfactory organs with a large sensory surface. The maintenance of a small repertoire of olfactory receptors may be necessary to support a high sensitivity for a small number of molecules. Indeed, it is likely that for a given number of olfactory neurons, there is a tradeoff, that is the higher the size of the olfactory repertoire, the lower the sensitivity for different molecules.

Secondly, the relative importance of the four olfactory receptor gene families is drastically different between cartilaginous and bony fishes. The VR2/OlfC family is by far the largest family in all cartilaginous fish examined, whereas in bony fish the OR family is far larger than the V2R/OlfC family. The ligands of cartilaginous fish V2R/OlfC receptors are unknown, but teleost V2R/OlfCs are activated by amino acids, which serve as food odors. Thus a large sensory surface together with a relatively small repertoire of V2R/OlfC receptors may enable extremely sensitive localization of prey.

### Olfactory gene expression patterns in catshark

Few vertebrate olfactory receptor repertoires have been analysed by comprehensive *in situ* hybridisation. However, the available information points to the expression of many if not most of olfactory receptor genes in mammalian and teleost fishes OSNs (Young et al. 2003; Alioto and Ngai 2006; Churcher et al. 2015; Yoon et al. 2015). This is in contrast to the situation we report here for the three minor shark receptor families. We could detect expression in OSNs only for one member per family for TAARs and V1R/ORAs, and for none of the 2 *tarl* genes and 8 o*r* genes examined. While we cannot rule out technical reasons or developmental differences (all tissues analysed stem from a similar stage, nearly adult juveniles), the rarity of expression for the TAAR and OR families could be related to the absence of ciliated OSNs in the shark olfactory organ (Theisen et al. 1986). Ciliated OSNs are the neuronal subpopulation which expresses ORs and TAARs in bony vertebrates (Hansen et al. 2004; Mombaerts 2004). Both families are very small and stable in all cartilaginous fishes examined, which would be consistent with a nonolfactory function in this taxon.

Expression profiling for several organs of two shark species showed broad expression for three ORs, with high expression levels observed only in non-olfactory organs (Hara et al. 2018). A comparison with lamprey suggests the absence of OR expression in OSNs to be a derived feature, since lampreys exhibit a moderately-sized OR family (Libants et al. 2009), show olfactory expression of ORs (Freitag et al. 1999) and do possess ciliated OSN expressing Golf, which is the G protein alpha subunit typically coupled to the OR and TAAR family (Frontini et al. 2003; Laframboise et al. 2007; Spehr and Munger 2009).

The TAAR family is absent in lamprey, which only possesses *taar-like (tarl)* genes (Grus and Zhang 2009; Hussain et al. 2009; Dieris et al. 2021). The absence of expression in OSNs for the two catshark *tarl* genes parallels the absence of OSN expression in teleost fish *tarl* and is in stark contrast to the expression of *tarl* genes in lamprey OSNs (Berghard and Dryer 1998; Dieris et al. 2021). This is consistent with the hypothesis that an olfactory function for *tarl* genes has been acquired independently in the jawless lineage, but not in cartilaginous or bony fishes (Dieris et al. 2021). Interestingly, the *taar* gene expressed in the catshark olfactory epithelium, *taar1a*, is the ortholog of a highly conserved *taar1* gene of bony vertebrates, which is non-olfactory in both tetrapods and teleosts (Liberles and Buck 2006; Hussain et al. 2009). Thus, catshark *taar1a* may have acquired olfactory function independently, possibly in microvillous receptor neurons, in contrast to *taar2-n* of bony fishes, which are expressed in ciliated neurons (Mombaerts 2004). We also found expression of only one *v1r/ora* gene in catshark OE, *ora2*. This is again different from the situation in zebrafish, where all *ora* genes show olfactory expression (Saraiva and Korsching 2007; Kowatschew et al., 2022). The cell type of the *ora2*-expressing neurons is unknown, but they could be microvillous neurons, which do express the related family of *v1r* genes in mammals (Mombaerts 2004).

In contrast to the three minor olfactory receptor families OR, V1R/ORA, and TAAR/TARL, each V2R/OlfC probe examined resulted in robust expression. Considering the cross-reactivity of several probes, we showed olfactory expression of up to 16 different V2R/OlfCs, a sizable proportion of the entire family. These results extend the olfactory function of the V2R family to the common ancestor of cartilaginous and bony fishes – the family is present with 1-2 genes in lamprey, but these are not expressed in the olfactory epithelium (Kowatschew and Korsching 2022). Notably, the *v2r1* probe showed a much higher density of labeled cells compared to the four probes cross-reacting with small subsets of genes. The *v2r1* gene is the ortholog of zebrafish *olfCc1* and mouse *vmn2r1-7*, which both have been shown to be co-expressed with many different individual *v2r/olfC* genes (Alioto and Ngai 2006; Silvotti et al. 2007; Ishii and Mombaerts 2011; DeMaria et al. 2013) and accordingly show a much higher density of expression compared to the individual genes. This suggests that the catshark *v2r1* could also serve as co-receptor (Ishii and Mombaerts 2011). The presence of a co-receptor is a special characteristic of the V2R/OlfC family, only paralleled by insect OR receptors, which are, however, evolutionarily unrelated to any vertebrate olfactory receptor (Yan et al. 2020).

Since the first discovery of olfactory receptor genes three decades ago, hundreds of expression studies have shown a common theme: individual receptor genes are expressed in sparsely distributed OSNs. Whenever these distributions have been examined more closely, they were observed to be different for different receptor genes, albeit often broadly overlapping. This ‘half-random’ feature has been described for mouse, rat, frog, zebrafish (Weth et al. 1996; Miyamichi et al. 2005; Syed et al. 2013; Zapiec and Mombaerts 2020) and recently also for lamprey (Kowatschew and Korsching 2022).

Here, we endeavored to examine whether this characteristic property of bony fishes and lamprey would also be present in cartilaginous fish. We do report that the basic principle of distinctly different spatial distributions for different olfactory receptor genes is present in the cartilaginous fish. This extends previous estimates derived from the comparison of tetrapods, teleost fishes and lamprey (Strotmann et al. 1996; Horowitz et al. 2014; Ahuja et al. 2018; Kowatschew and Korsching 2022).

It is unclear whether such differences in spatial expression patterns might have functional meaning. The presence of secondary lamellae in the catshark olfactory organ might serve just to increase the surface area of the sensory surface (Ferrando et al. 2019), but differences in access of odorants to primary vs. secondary lamella areas cannot be excluded since existing studies of hydrodynamic properties focus solely on primary lamellae (see e.g. Cox 2008). Alternatively, differences in radial and primary/secondary lamella distribution could result as consequence of the developmental mechanisms guiding the olfactory receptor expression (*cf*. Bayramli et al. 2017).

Taken together we have shown for the first time the cellular expression of olfactory receptors in a cartilaginous fish. The expression is dominated by *v2r/olfC* genes, with minor contributions from a *v1r/ora* and a *taar* gene. The spatial expression patterns of different receptor genes are characteristically different, both for a topological parameter shared with bony fishes (radial parameter) and for a peculiar property of all cartilaginous fishes (secondary lamellae). Thus the principle of non-random spatial organization is shared between jawless, cartilaginous, and bony vertebrates, suggesting similarly broad presence of the underlying molecular mechanisms.

## Conclusion

Comparative studies showed that olfactory receptors belonging to the OR, TAAR and V1R/ORA families were co-opted early during the evolution of vertebrates and were present in the last common ancestor of extant vertebrates. Although V2R/OlfC receptors were also present, there were co-opted as olfactory receptors later, after the separation of jawless fishes and jawed vertebrates. In bony fishes V2R/OlfCs are less abundant than ORs and TAARs, but in cartilaginous fishes V2R/OlfCs constitute the essential component of the entire olfactory receptor repertoire. Why such a difference? Conceivably this could be an indirect effect of the loss of the ciliated subtype of olfactory sensory neurons which are expected to express ORs and TAARs. Whether the dearth of ORs and TAARs amounts to a restriction in the odor space accessible to cartilaginous fish remains to be seen. The correlation between the number of OR, TAAR and V2R/OlfC receptors in ray-finned fishes might suggest an initial functional overlap between these receptor families which in tetrapods have divergently evolved to detect volatile substances in the main olfactory epithelium and non-volatile substances in the vomeronasal organ. Overall, cartilaginous and jawless fish have a small olfactory receptor gene repertoire compared to bony fishes, despite having very well-developed olfactory organs. These divergent evolutionary trajectories could result from different tradeoffs between sensitivity and odor discrimination, with cartilaginous fish maximizing sensitivity of odor detection.

Within cartilaginous fishes there are notable differences in the number of olfactory receptors, which remain to be understood. For example, the thorny skate *Amblyraja radiata* may have less than ten OlfC receptors, whereas the chimaera *C. milii* has more than 40 OlfC receptors, although they consume relatively similar diet. Further behavioral and functional studies will be necessary to better understand this issue.

## Materials and Methods

### Genome dataset and species phylogeny

Seventeen chondrichthyan genome assemblies, corresponding to thirteen species were downloaded from NCBI: *Amblyraja radiata* (GCF_010909765.2), *Callorhinchus milii* (GCF_018977255.1 ; GCA_000165045.2), *Carcharodon carcharias* (GCF_017639515.1 ; GCA_003604245.1), *Chiloscyllium plagiosum* (GCF_004010195.1), *Chiloscyllium punctatum* (GCA_003427335.1), *Hemiscyllium ocellatum* (GCA_020745735.1), *Hydrolagus affinis* (GCA_012026655.1), *Leucoraja erinacea* (GCA_000238235.1), *Pristis pectinata* (GCA_009764475.2), *Rhincodon typus* (GCA_001642345.3 ; GCA_013626285.1 ; GCA_013626285.1), *Scyliorhinus canicula* (GCA_902713615.2), *Scyliorhinus torazame* (GCA_003427355.1), *Stegostoma fasciatum* (GCA_022316705.1).

The completeness of these genomes was assessed with BUSCO v5.1.2 using the vertebrata_odb10 database (Manni et al., 2021). For species with multiple genome assemblies, we retained only the one for which BUSCO retrieved the highest number of complete genes (supplementary table S1, Supplementary Material online). *Leucoraja erinacea* was removed from the phylogenetic analysis described below, as only 7% of BUSCO genes could be retrieved complete from its genome assembly (supplementary fig. S6, Supplementary Material online). We then extracted protein sequences of 1068 BUSCO genes that were retrieved in common in single copy in the best assemblies for each species, and align these sequences individually using MAFFT (auto, v7.407) (Katoh and Standley 2013). These alignments were trimmed using trimal v1.4.1 (with the option -automated1) (Capella-Gutiérrez et al., 2009) and concatenated using AMAS (Borowiec 2016). A maximum likelihood phylogeny was then computed with IQ-TREE v2.2.0 (Minh et al., 2020) and the best model for each partition was assessed with ModelFinder (option -m MFP+MERGE) (Kalyaanamoorthy et al., 2017). The least square dating method implemented in IQ-TREE was used to build a time-calibrated phylogeny from the inferred tree topology (with the options --date-tip 0 --date-ci 100). Five calibration dates retrieved on Timetree.org were used (supplementary fig. S5, Supplementary Material online) (Kumar et al., 2022).

### Olfactory receptor gene mining

*or, taar, v1r/ora* and *v2r/olfC* genes were mined in the most complete genome assemblies of each species, except for *L. erinacea* and *H. affinis* for which only 7% respectively 54% of BUSCO genes could be retrieved complete. Thus an accurate estimation of the number of olfactory genes was not possible in these two species. The naming for *S. canicula* olfactory receptor genes was based on (Sharma et al., 2019), for changes/additions see (supplementary table S2, Supplementary Material online).

Gene mining was performed following methods described by (Policarpo et al. 2022). Briefly, TBLASTN searches (e-value < 1e-10) were performed against genome assemblies using known olfactory receptors belonging to the four families in other vertebrate species as queries. Non-overlapping hit regions were extracted and extended using SAMTools and genes were predicted on those regions using EXONERATE (options : --model protein2genome – minintron 50 –maxintron 20000). We verified that predicted genes were true olfactory receptors with a BLASTX against a custom database of olfactory, taste and other G-protein-coupled receptors and with phylogenetic trees, retaining only sequences that clustered with known olfactory receptors. Retrieved sequences were then classified into four categories: 1) “complete” if a complete coding sequence was retrieved; 2) “pseudogene” if the coding sequence contained at least one loss-of-function mutation (a premature stop codon or a frameshift); 3) “incomplete” if the gene was found incomplete and with no loss-of-function mutation; 4) “edge” if the gene was found incomplete and near a contig or scaffold border, which are most likely assembly artifacts. For nucleotide sequences for all validated olfactory receptor genes see (supplementary file S2, Supplementary Material online).

Protein sequences of complete genes obtained in the previous step, as well as outgroup protein sequences (see supplementary file S2, Supplementary Material online) were aligned using MAFFT v7.467 and maximum likelihood phylogenies were computed with IQ-TREE 2.0 with the best model found by ModelFinder. Branch supports were obtained with 1000 ultrafast bootstraps.

### Patterns of gene birth and death

A gene tree – species tree reconciliation method was used to infer the number of gene duplications and gene losses in every branch of the species tree. We first collapsed nodes with low bootstrap values (<90%) in gene phylogenies for the four olfactory receptor families using the R package ape v5.0 (Paradis and Schliep 2019). Treerecs was then used to find the best root and reconcile gene trees with the species tree, with default parameters (Comte et al 2020).

### Tissue preparation

Paraformaldehyde-fixed whole olfactory organs from 2 nearly adult juvenile catsharks were kindly provided by Sylvie Mazan and Ronan Lagadec. Organs were stored in methanol at - 80°C. Before using tissues were rehydrated in decreasing concentrations (75%, 50%, 25%) of methanol and rinsed thrice in PBS, followed by equilibration in 15 % saccharose in PBS at 4°C until they sank. They were then equilibrated in 30% saccharose in PBS and embedded in Tissue Tek.

### mRNA Isolation, cDNA Synthesis and RT-PCR

Total mRNA was extracted using the easy-spin Total RNA Extraction Kit (iNtRON Biotechnology) resulting in highly concentrated mRNA (OD260 > 0.9). Complementary DNA (cDNA) was synthesized from olfactory organ using SuperScript II Reverse Transcriptase (Invitrogen, No. 18064022). cDNA concentration was determined with a NanoDrop TM photometer and samples stored at -20°C. Forward (Fwd) and reverse (Rev) primers for each gene were chosen to result in fragment lengths between 350 and 560 bp (supplementary table S3, Supplementary Material online). For RT-PCR, annealing temperatures between 55 °C and 58 °C were used. RNA probes for in situ hybridisation were generated from PCR product by adding T3 promoter sequence (ATTAACCCTCACTAAAGG) 5′ to the Rev primer (supplementary table S3, Supplementary Material online).

### *In situ* hybridisation, stand-alone and combined with immunohistochemistry

Transverse cryostat sections of 10 μm were obtained (Leica CM1900) and dried, according to the cutting plane shown in fig. 1a. *In situ* hybridisation was done as described (Ahuja et al. 2018). Immunofluorescence was done as described (Ahuja et al. 2014). Primary antibodies used in the Immunofluorescence are mouse anti-PCNA (1:200, Merck) and mouse anti-HuC (1:200, Invitrogen) anti-Go (K-20) antibody (rabbit IgG; 1:50; sc-387, Santa Cruz Biotechnology). Secondary antibodies used were goat anti-rabbit IgG conjugated to Alexa Fluor 488 (A21206, Invitrogen) or Alexa Fluor 594 (A11012, Invitrogen). 4′,6-diamidino-2-phenylindole (DAPI) was used as counterstain for fluorescent detection. Micrographs were taken using a Keyence BZ-9000 fluorescence microscope, and absence of crosstalk between channels was confirmed. Double labeling was done by combining ISH and IHC as described (von Twickel et al. 2019).

### Quantification and statistical evaluation

Image analysis - lengths and distances measurements and cell counts - was performed using ImageJ (Schneider et al. 2012). The normalized position of labeled cells along the length of a lamella was determined by dividing the distance from the apex (toward the center of the olfactory rosette) by the total length of the lamella (from the base, which is attached to the peripheral connective capsule to the apex). For neurons situated in secondary lamellae a projection onto the corresponding primary lamella was used.

To determine the density, labeled neurons were counted in up to nearly 400 mm of lamellar length to achieve counts of at least one hundred cells per gene. For the most frequently expressed gene, *v2r1*, nearly 100 mm of lamellar length were evaluated, which contained over 400 labeled neurons.

Radial distributions are shown as empirical cumulative distribution function (ECDF) (Feller 1966, Wilk and Gnanadesikan 1968). To estimate whether two distributions were significantly different, we performed Kolmogorov-Smirnov tests as implemented in R version 4.1.0 (Marsaglia et al. 2003).

## Supporting information

Supplementary Figure S1. Olfactory receptor gene trees

Supplementary Figure S2. RT-PCR shows expression of all but one olfactory receptor examined

Supplementary Figure S3. Phylogenetic position of genes analysed for expression

Supplementary Figure S4. Expression of a taar and an ora gene in sparse OSN

Supplementary Figure S5. Complete Species tree

Supplementary Figure S6. BUSCO results

Supplementary Table S1. Genomes analysed and number of olfactory receptor genes

Supplementary Table S2. Correspondances and name changes for small-spotted catshark olfactory receptors published in Sharma et al 2019

Supplementary Table S3. Primer sequences employed for RT-PCR and in situ hybridisation

Supplementary file S1. Legends for supplementary files

Supplementary file S2. Nucleotide Sequences All OlfR genes

## Data availability

Concatenated gene alignment of the 1068 BUSCO genes as well as the calibrations points used and the species tree in nexus format can be found in Figshare: https://doi.org/10.6084/m9.figshare.21507615.v1

## Supplementary Material

Supplementary data are available at Molecular Biology and Evolution.

## Acknowledgements

We are grateful to Sylvie Mazan (Genoscope-Centre National de Séquençage, Evry, France) and Ronan Lagadec (Observatoire Océanologique de Banyuls-sur-mer UMR7232 BIOM - CNRS/UPMC, France) for kindly providing the catshark olfactory tissues used in this work. We are thankful to Daniel Kowatschew for expert technical advice. This work was supported by the German Science foundation (grant KO1046/12-1 to S.I.K. and grant SH1768/1-1 to K.S.).

## Author contributions

S.I.K., A.S. and K.S. conceived the study; A.S. and K.S. designed the experiments; A.S., K.S., S.F. and M.P. performed the experiments and conducted the analysis. S.I.K., K.S., S.F., M.P. and D.C. prepared the figures and wrote the manuscript. All authors read and approved the manuscript.

