## Supplementary Figure S1. Olfactory receptor gene trees for "Ancient and nonuniform loss of olfactory receptor expression renders the shark nose a *de facto* vomeronasal organ"

## A - OR

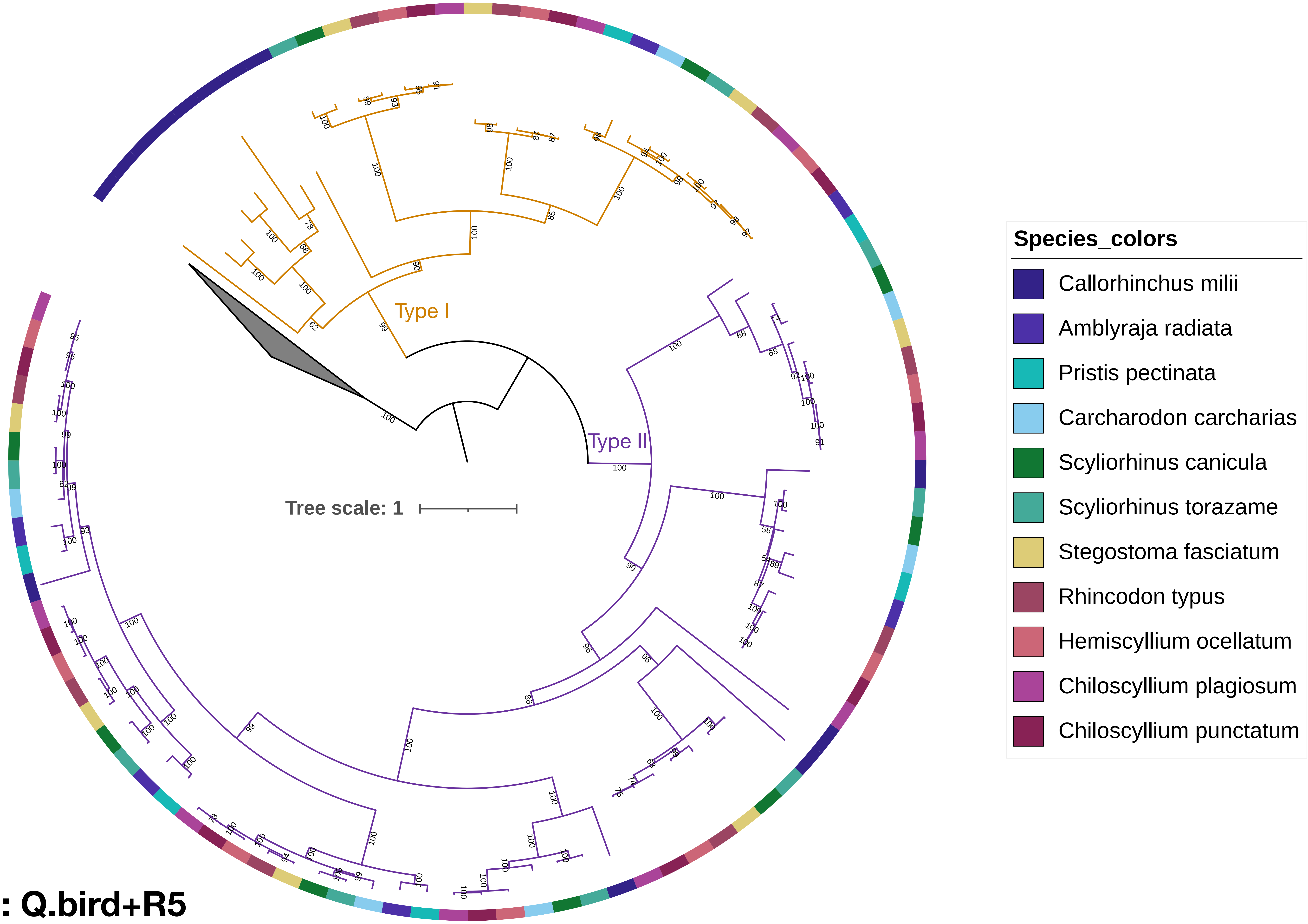


**Substitution model : Q.bird+R5**

### B - ORA

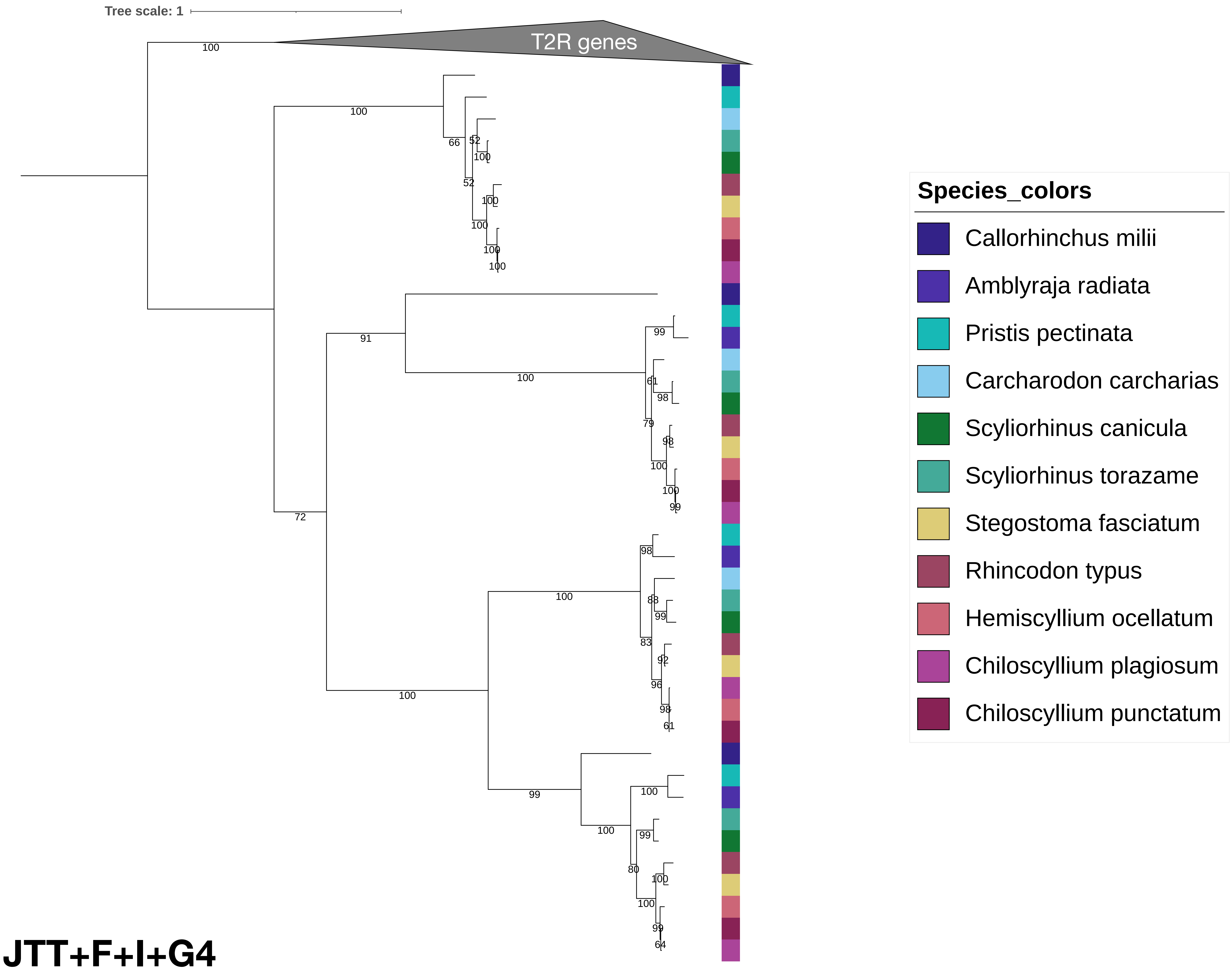


#### Substitution model : JTT+F+I+G4

### C - TAAR + TARL

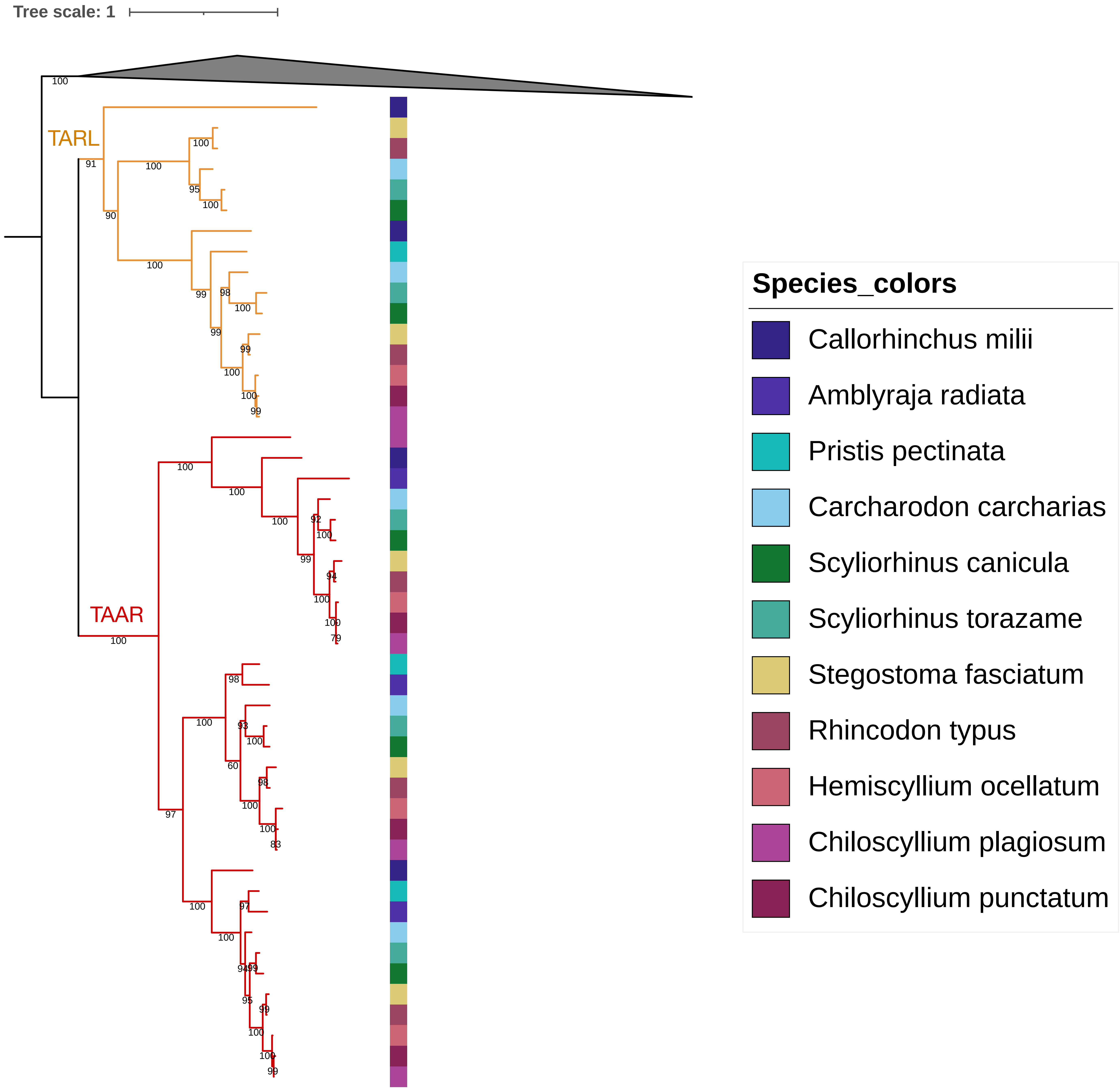


#### Substitution model : Q.plant+R5

D - V2R

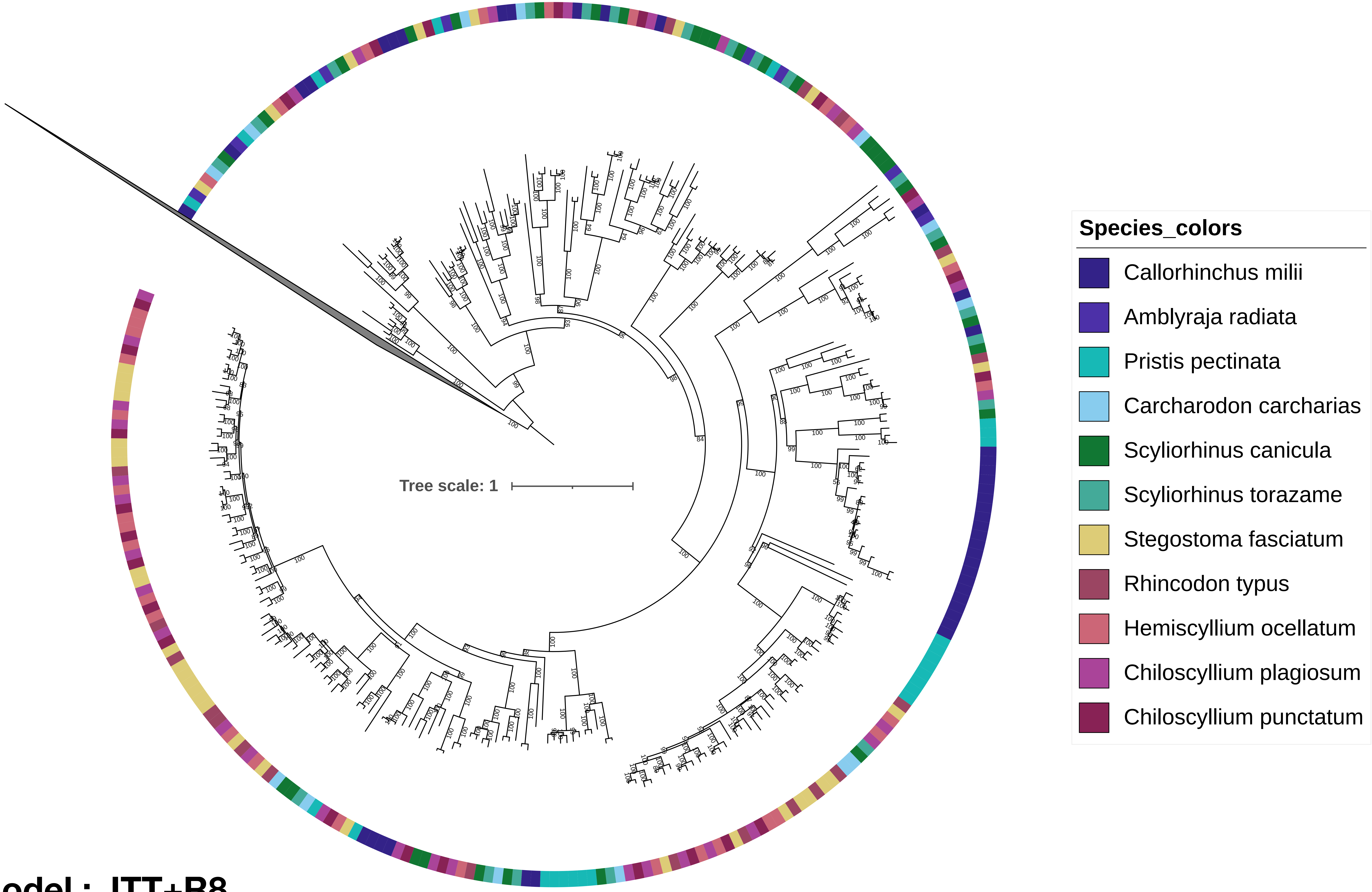


Substitution model : JTT+R8
