## Supplementary figures and images for "Ancient and nonuniform loss of olfactory receptor expression renders the shark nose a *de facto* vomeronasal organ"

### Supplementary Figure S2. RT-PCR shows expression of all but one olfactory receptor examined

#### RT-PCR: expression of olfactory receptors from 4 families in the OE

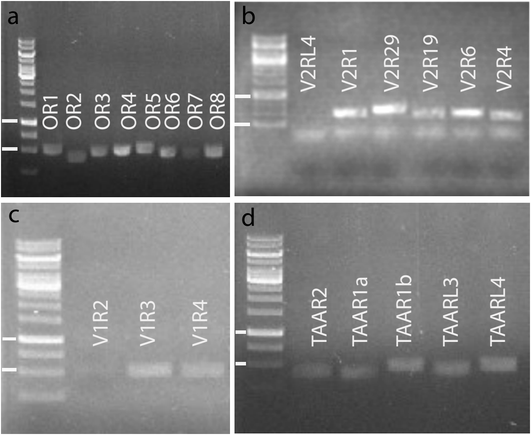

### Supplementary Figure S3. Phylogenetic position of genes analysed for expression

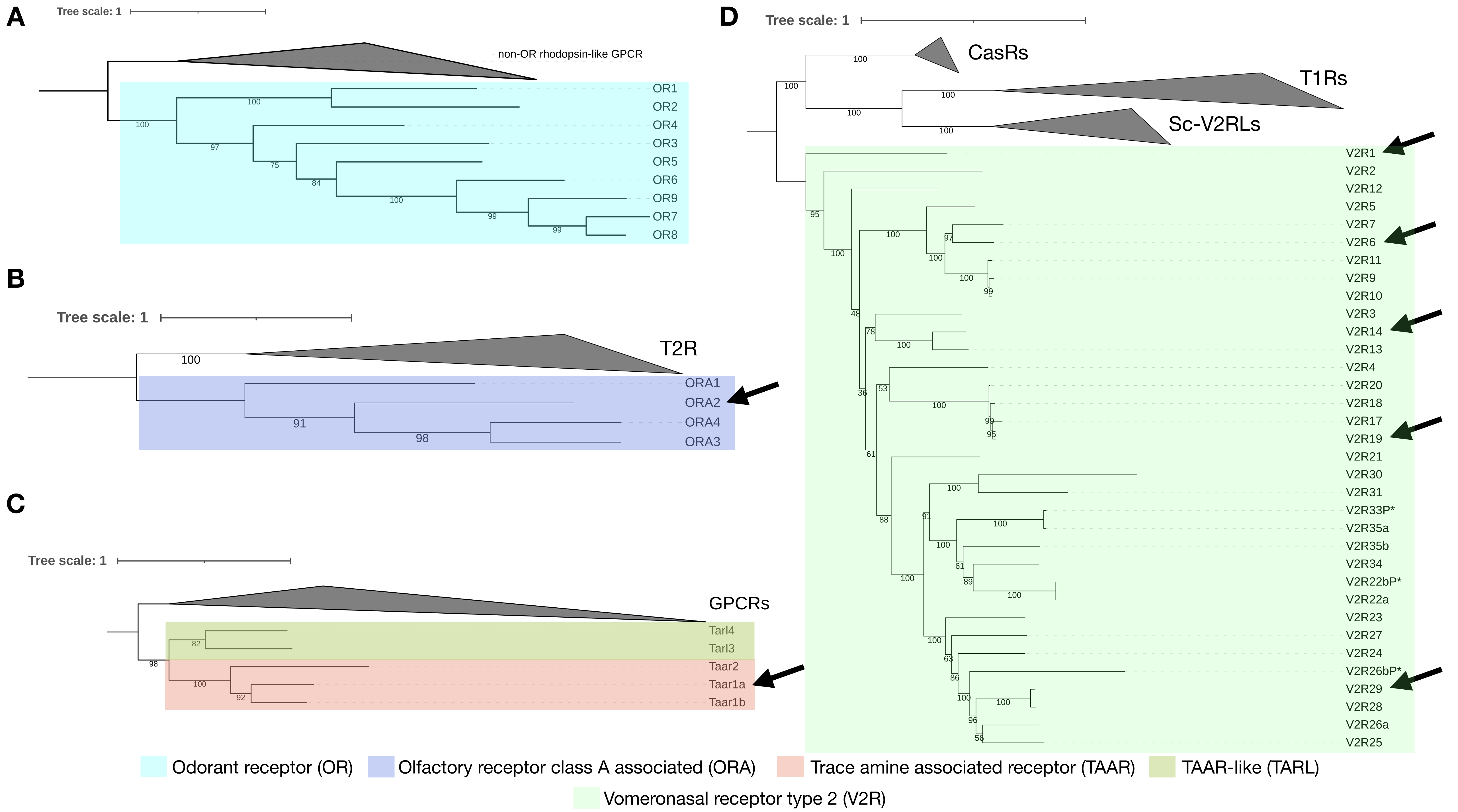

### Supplementary Figure S4. Expression of a taar and an ora gene in sparse OSN

#### Expression of ORA2 and TAAR1a in OE

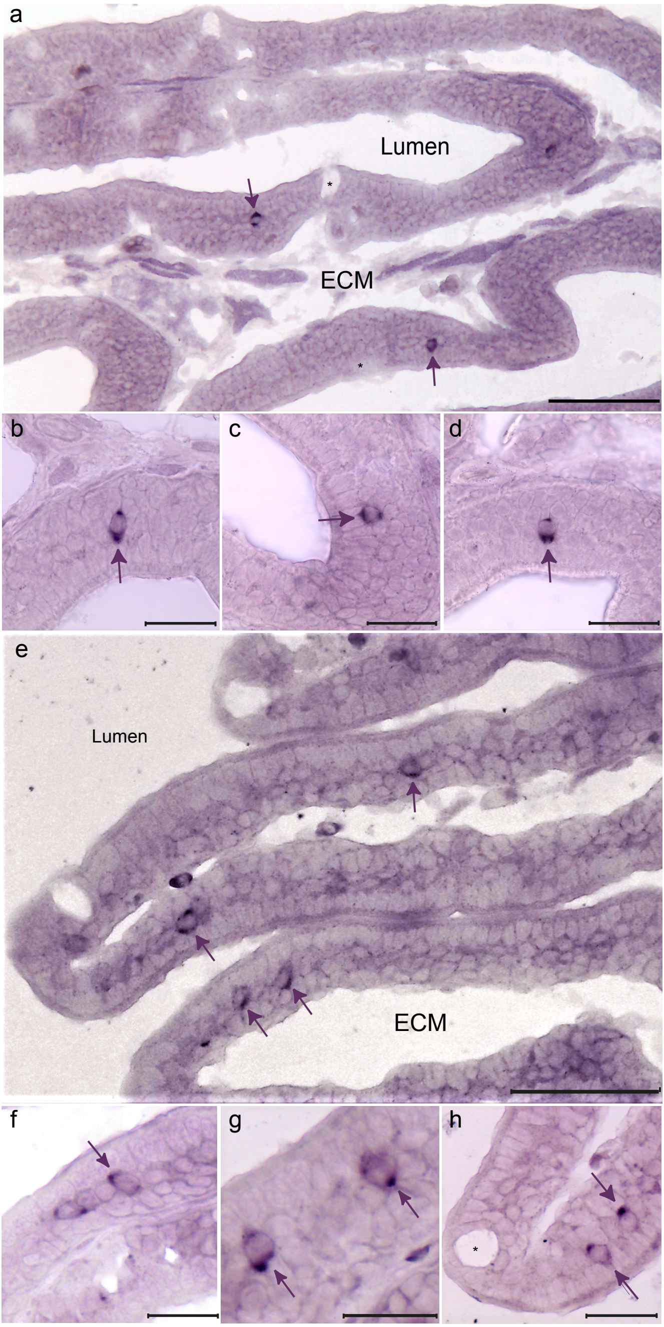

### Supplementary Figure S5. Complete Species tree

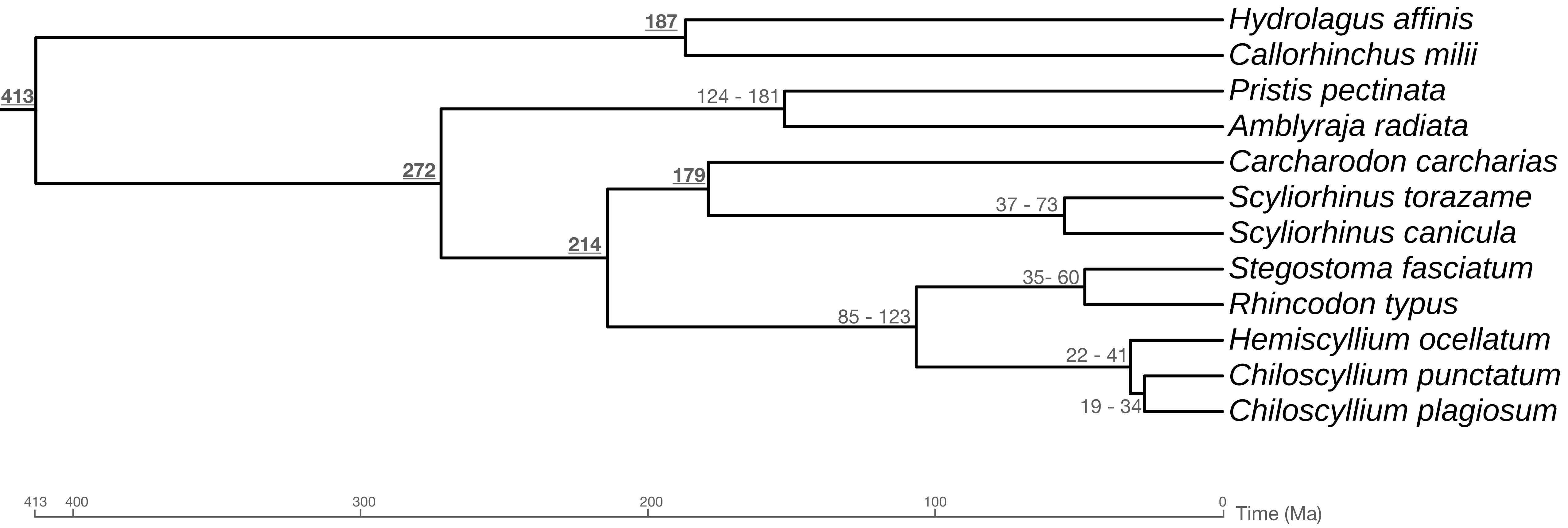

### Supplementary Figure S6. BUSCO results

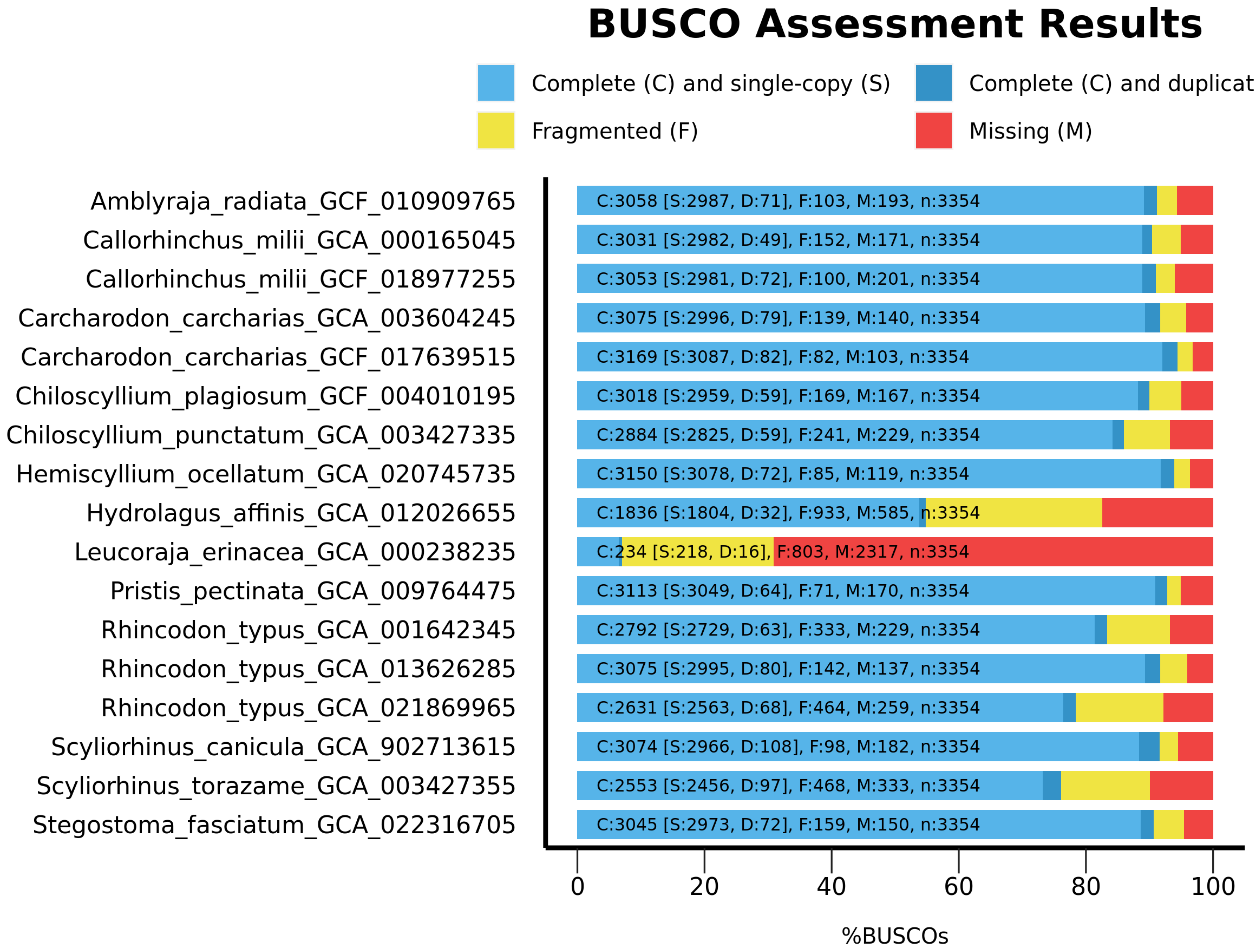
