## Supplementary file S1. Legends for supplementary files for "Ancient and nonuniform loss of olfactory receptor expression renders the shark nose a *de facto* vomeronasal organ"

Legends for Supplementary Figures and Tables

Supplementary File S1 Legends for Supplementary Figures and Tables

Supplementary File S2 Nucleotide sequences of olfactory receptor genes of cartilaginous fish. Full length and incomplete sequences identified in this study are given as zip archive.

Supplementary Figure S1 Olfactory receptor gene trees. Phylogenies of olfactory receptor genes retrieved in eleven chondrichthyes (A- or ; B- v1r/ora ; C-taar ; D- v2r/olfC). Numbers at internal nodes indicate the support values computed with 1000 UltraFast bootsraps. The best model found by ModelFinder for each family is indicated in the lower right corner. Colorstips indicate to which species each gene belongs to. Or genes were rooted using six non-or rhodopsin-like GPCR genes retrieved from Niimura et al. 2009 (NP 000670.1; NP 037477.1 ; NP 005292.2 ; NP 001287.2 ; NP 001044.1 ; NP 000729.2). ORA genes were rooted with twelve Tas2r genes of vertebrate species retrieved on NCBI (XP 021021573.1 ; BAM38240.1 ; NP 076407.1 ; NP 001018341.1 ; NP 001034919.1 ; XP 006640475.1 ; XP 022074252.1 ; XP 022623854.1 ; XP 026117515.1 ; XP 026126163.1 ; XP 026125823.1 ; XP 018587155.1). TAAR genes were rooted with seventeen non-TAAR rhodopsin-like GPCR genes retrieved on NCBI (BC162681.1 ; BC163374.1 ; BC163360.1 ; XP 003452776.1 ; NP 001037811.1 ; AB481209.1 ; BC163344.1 ; XP 003454257.1 ; XP 003452229.1 ; XP 003447064.1 ; XP 003441904.1 ; CBN80867.1 ; XP 003454619.1 ; BC163414.1 ; EU729324.1 ; BC163924.1 ; BC163917.1). V2R genes were rooted with six CasR and eight Tas1R genes of vertebrates species retrieved on NCBI (NP 001338594.1 ; AAI62412.1 ; NP 001077325.1 ; NP 001034920.1 ; ABK81648.1 ; NP 114073.1 ; NP 114079.1 ; NP 114078.1 ; AAB46873.1 ; NP 038831.2 ; NP 776427.1 ; NP 001296567.1 ; XP 005654095.1 ; NP 689418.2).

Supplementary Figure S2 RT-PCR shows expression of all but one olfactory receptor examined. RT-PCR showed presence of the olfactory receptor repertoire in the olfactory epithelium of catshark. (a-d) or, v2r, ora/v1r, and taar respectively. White dashes on ladder indicate 1000bp and 500 bp. Annealing temperatures between 55 °C and 58 °C were used with cDNA as template resulting in fragment lengths varying from 350 to 560 bp.

Supplementary Figure S3 Phylogenetic position of genes analysed for expression. Results of expression analysis are mapped onto a maximum likelihood tree of the entire olfactory receptor repertoire of catshark (35 v2r, 9 or, 6 taar+tarl, 3 ora genes – note that genes named ora5-711 place within the rhodopsin clade, when this clade is present in the outgroup). Branch support at the basal node of each receptor family is given in %. Each phylogenetic tree was computed with IQ-TREE with 1000 ultrafast bootstraps and the best model found by ModelFinder. OR genes were rooted using six non-OR rhodopsin-like GPCR genes retrieved from Niimura et al. 2009 (NP 000670.1; NP 037477.1 ; NP 005292.2 ; NP 001287.2 ; NP 001044.1 ; NP 000729.2). ORA genes were rooted with twelve Tas2r genes of vertebrate species retrieved on NCBI (XP 021021573.1 ; BAM38240.1 ; NP 076407.1 ; NP 001018341.1 ; NP 001034919.1 ; XP 006640475.1 ; XP 022074252.1 ; XP 022623854.1 ; XP 026117515.1 ; XP 026126163.1 ; XP 026125823.1 ; XP 018587155.1). TAAR genes were rooted with seventeen non-TAAR rhodopsin-like GPCR genes retrieved on NCBI (BC162681.1 ; BC163374.1 ; BC163360.1 ; XP 003452776.1 ; NP 001037811.1 ; AB481209.1 ; BC163344.1 ; XP 003454257.1 ; XP 003452229.1 ; XP 003447064.1 ; XP 003441904.1 ; CBN80867.1 ; XP 003454619.1 ; BC163414.1 ; EU729324.1 ; BC163924.1 ; BC163917.1). V2R genes were rooted with six CasR and eight Tas1R genes of vertebrates species retrieved on NCBI (NP 001338594.1 ; AAI62412.1 ; NP 001077325.1 ; NP 001034920.1 ; ABK81648.1 ; NP 114073.1 ; NP 114079.1 ; NP 114078.1 ; AAB46873.1 ; NP 038831.2 ; NP 776427.1 ; NP 001296567.1 ; XP 005654095.1 ; NP 689418.2) and with five V2RL sequences of catshark retrieved in Sharma et al. 2019. Catshark gene were named according to Sharma et al. 2019 with few changes for additional genes or corrected gene annotations made in this study (See supplementary table 2, Supplementary Material online, Correspondances and name changes Sc Olfactory receptors Sharma et al 2019.xlsx). Pseudogene names end with P* while truncated gene names end with T*. All other genes were found complete. Arrows point to genes whose expression is observable by *in situ* hybridization. Each receptor family is overlayed by colored rectangle, color code as indicated.

Supplementary Figure S4 Expression of a *taar* and an *ora* gene in sparse OSN. Horizontal cryostat sections of catshark olfactory epithelium were hybridized with probes for ora2 (a-d) and taar1a (e-h). Both probes show expression in small subsets of scattered OSNs situated in the middle layer of the sensory epithelium, along both primary and secondary lamellae. ECM, extracellular matrix. Scale bars, 100 μm for panel a,e) and 40 μm for panels b-d and f-h)

Supplementary Figure S5 Complete Species tree The species tree topology was inferred by maximum likelihood using 1068 concatenated BUSCO genes and node ages were inferred using the least square dating method. The tree is the same as in figure 2 but with one more species (*Hydrolagus affinis*) that did not have a high enough BUSCO score to correctly infer the number of olfactory receptors.

Supplementary Figure S6 BUSCO results. BUSCO assessment result for 17 chondrichthyes genome assemblies. BUSCO (v5.1.2) was run with default parameters using the vertebrata odb10 database.

Supplementary Table S1 Genomes analysed and number of olfactory receptor genes.xlsx. Sheet 1 : NCBI Assembly accession of retained genome for each species. Sheet 2, 3,4,5 : Number of ora, or, taar and v2r/olfC genes retrieved in each categories for the 11 species.

Supplementary Table S2 Correspondances and name changes for small-spotted catshark olfactory receptors published in Sharma et al 2019. Correspondance of olfactory receptor genes retrieved in the small-spotted catshark genome assembly between this study and the study of Sharma et al. 2019, which used a preliminary genome draft. The first column indicates the gene family, the second column the names of the genes in this study (their sequences are given in the Supplementary File S2). The third column indicates the corresponding gene name in Sharma et al. 2019 as well as the type of change that was made: "No changes" means that the exact same sequence was found and the gene name was kept; "Merge" indicates that several incomplete genes in Sharma et al. 2019 were retrieved as a single complete gene in this study, leading to a change in the gene name. Four genes retrieved here were "Not found" in Sharma et al. 2019. These genes were given new names.

Supplementary Table S3 Primer sequences employed for RT-PCR and *in situ* hybridisation. Primers were used to perform PCR with cDNA from catshark olfactory epithelium. Forward and reverse primer sequences of expressed olfactory genes are listed. The T3 promoter sequence (ATTAACCCTCACTAAAGG) was added 5′ to the reverse primer for generation of RNA probes. Number of bp denotes the length of the PCR product used for generation of the RNA probes.
